# Hidden conformations differentiate day and night in a circadian pacemaker

**DOI:** 10.1101/2021.09.14.460370

**Authors:** Jeffrey A. Swan, Colby R. Sandate, Archana G. Chavan, Alfred M. Freeberg, Diana Etwaru, Dustin C. Ernst, Joseph G. Palacios, Susan S. Golden, Andy LiWang, Gabriel C. Lander, Carrie L. Partch

**Affiliations:** Department of Chemistry & Biochemistry, University of California, Santa Cruz, CA 95064; Department of Integrative Structural and Computational Biology, The Scripps Research Institute, La Jolla, CA 92131; Department of Chemistry and Biochemistry, University of California, Merced, CA 95343; Division of Biological Sciences, University of California, San Diego, La Jolla, CA 92093; Center for Circadian Biology, University of California, San Diego, La Jolla, CA 92093

**Author notes:** These authors contributed equally to this work (Jeffrey A. Swan, Colby R. Sandate).

## Abstract

The AAA+ protein KaiC is the central pacemaker for cyanobacterial circadian rhythms. Composed of two hexameric rings with tightly coupled activities, KaiC undergoes changes in autophosphorylation on its C-terminal (CII) domain that restrict binding of of clock proteins on its N-terminal (CI) domain to the evening. Here, we use cryo-electron microscopy to investigate how daytime and nighttime states of CII regulate KaiB binding to CI. We find that the CII hexamer is destabilized during the day but takes on a rigidified C_2_-symmetric state at night,concomitant with ring-ring compression. Residues at the CI-CII interface are required for phospho-dependent KaiB association, coupling ATPase activity on CI to cooperative KaiB recruitment. Together these studies reveal how daily changes in KaiC phosphorylation regulate cyanobacterial circadian rhythms.

**One-Sentence Summary:** Cryo-EM structures of KaiC in its day and night states reveal the structural basis for assembly of clock regulatory complexes.

## Main Text

Cyanobacteria possess an internal circadian clock that temporally aligns gene expression with the solar day to maximize photosynthetic output and coordinate integrated metabolic processes (*1-3*). The most basic manifestation of this timekeeping function is a cyclic pattern of autophosphorylation in the C-terminal (CII) domain of the hexameric clock protein KaiC (*4*) that results in the following sequence of post-translational modifications at residues S431 and T432: S/T ⍰ S/pT ⍰ pS/pT ⍰ pS/T (*5, 6*). During the day, autophosphorylation is stimulated by another clock protein, KaiA; and at night, compression of the two KaiC rings opens the N-terminal (CI) ring and exposes binding sites that allow a third clock protein, KaiB, to interact with KaiC and sequester KaiA (*7*). Without KaiA bound to CII, KaiC begins to autodephosphorylate until KaiB and KaiA are released from CI, thus completing the negative feedback loop. Both *in vitro* and *in vivo* studies have linked the S/T and S/pT phosphostates of KaiC with daytime, while the pS/pT and pS/T states are associated with the KaiB-associated nighttime state of KaiC (*5, 6*). The structural basis for formation of the nighttime KaiABC complex is understood (*8, 9*), however no direct mechanistic link between CII phosphorylation and regulation of KaiB binding has yet been identified.

KaiC can be trapped in its daytime or nighttime state using phosphomimetic substitutions at phase-defining residues (*6*), referred to herein as KaiC-AE for the daytime variant (S431A-T432E) and KaiC-EA for the nighttime variant (S431E-T432A, see **Table S1**). Despite stark differences in clock protein association, previous crystallographic studies of the phosphomimetic forms did not reveal structural changes that might explain their distinct biochemical properties (*10*). Nevertheless, solution biophysical studies have demonstrated overall changes in the shape of KaiC throughout the phosphocycle (*11, 12*) as well as changes in intra- and inter-subunit interactions of the adenosine triphosphatase (ATPase) domains (*13*). Here, we present cryo-electron microscopy (cryo-EM) structures of daytime and nighttime KaiC that reveal new conformations demonstrating how temporal information encoded in CII phosphorylation is transmitted through dynamic structural features to regulate the cyanobacterial circadian clock.

To establish a baseline for KaiB discrimination between the phosphomimetics, we first compared KaiB affinity in the daytime and nighttime-trapped forms of KaiC (**Figs. 1A, S1A**). We observed ≥30-fold tighter binding of KaiB to nighttime KaiC than the daytime variant, recapitulating previously observed day/night distinction between the KaiC-AE and KaiC-EA phosphomimetics (*6*). Next, we subjected KaiC-EA and KaiC-AE to comparison by cryo-EM in the presence of a saturating concentration of adenosine triphosphate (ATP) to identify structural differences between the phosphomimetic variants in solution.

**Fig. 1.**
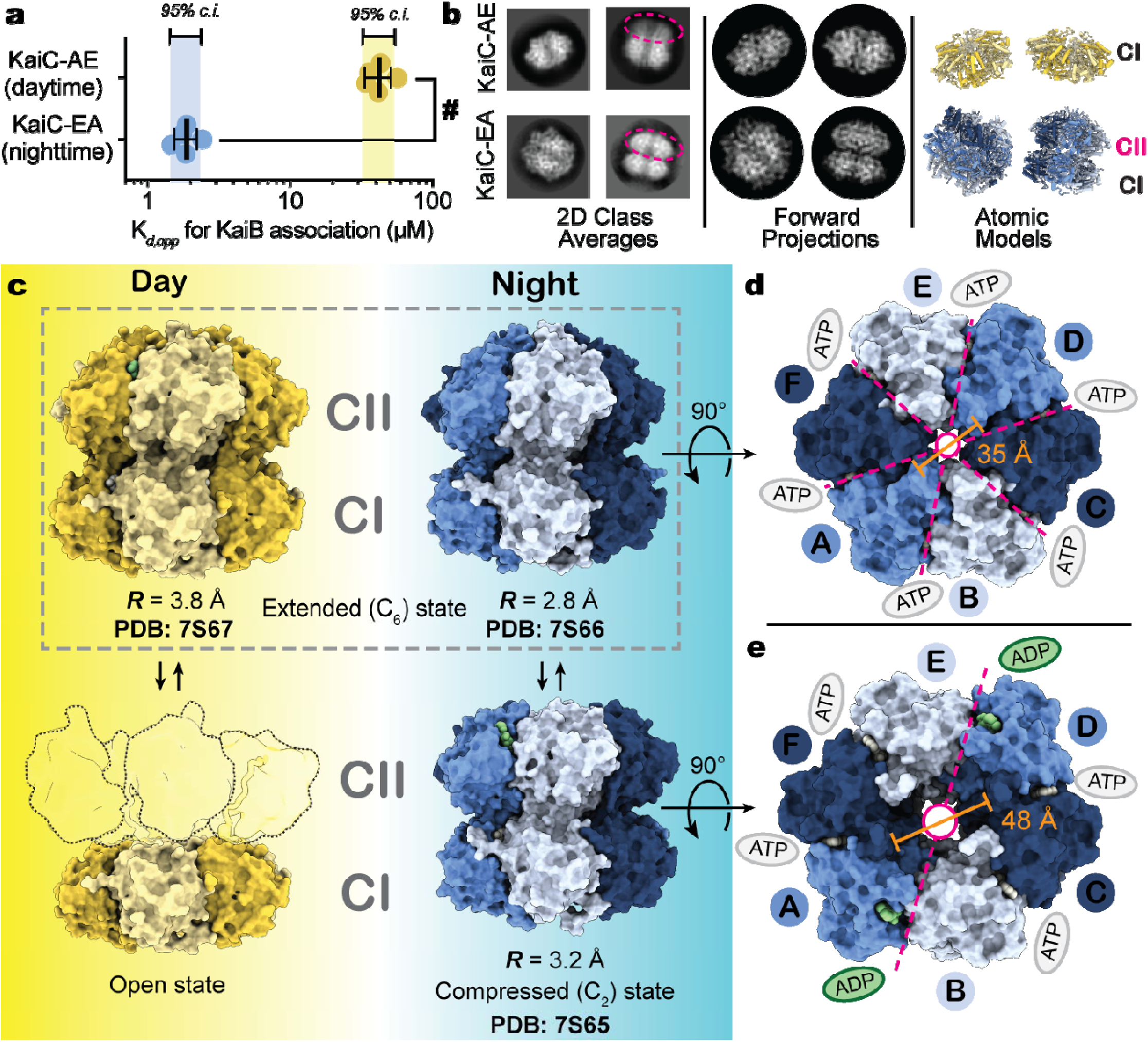
Daytime and nighttime phosphomimetics confer distinct biochemical activities and global conformations to KaiC. **a**) Apparent binding constants (K_*d,app*_) for KaiB association with daytime or nighttime KaiC variants in units of protomer concentration. A hashtag (**#**) represents *P* < 0.0001 from an unpaired parametric t-test. Lighter colored regions represent the 95% confidence interval for n = 5 measurements. **b**) Reference-free 2-dimensional class averages from electron micrographs of KaiC phosphomimetics. Dashed ovals indicate the CII rings, where visible, as inferred from forward projections obtained using the atomic models shown. **c**) Space-filling depictions of alternate daytime and nighttime KaiC conformations observed by cryo-EM. The ‘open’ daytime state is depicted with outlines to represent the destabilized CII protomers as rigid bodies flexibly tethered to the visible CI hexamer. **d-e**) Axial view of the extended (**d**) and compressed (**e**) nighttime KaiC structures, viewed from the side of the CII. Dashed lines indicate the axes of symmetry with colored ovals denoting nucleotide state in the CII ring. Pore diameters are reported as measured between C_α_ atoms of V433 on A and D protomers.

In daytime KaiC, we did not observe well-defined cryo-EM density for the CII ring (**Fig. 1B**), consistent with previous solution studies where daytime KaiC has exhibited an ‘open’ or destabilized CII hexamer (*13, 14*). Since flexible domains are often susceptible to damaging interactions with the hydrophobic air-water interface during sample preparation, we also structurally characterized daytime KaiC in the presence of perfluorinated fos-choline, which is known to limit air-water-interface interactions. This resulted in the same C_6_-symmetric ‘extended’ conformation previously observed by crystallography in the presence of non-hydrolyzable ATP analogue (*10*) (**Figs. S2, 1C, PDB: 7S67**), this time resolved to ∼3.8 Å. However, in our structure, adenosine diphosphate (ADP) was situated at the CII-CII interfaces, suggesting that CII nucleotide state does not influence global conformation in the ground state of daytime KaiC. Notably, the daytime and nighttime KaiC phosphomimetics maintained functional discrimination for KaiB in the presence of fos-choline (**Fig S3**), consistent with the idea that these open and extended conformations are both present in the solution structural ensemble of daytime KaiC.

In the nighttime phosphomimetic, single particle analysis resulted in two distinct maps with both the CI and CII rings resolved (**Figs. S4, 1B**). Once again, we observed the C_6_-symmetric ‘extended’ conformation, this time at ∼2.9 Å resolution (**PDB: 7S66**), as well as a population of particles exhibiting a novel C_2_-symmetric structure, resolved to ∼3.2 Å (**PDB: 7S65**). This subpopulation of KaiC hexamers has a widened central pore (**Fig. 1D-E**), coupled with accumulation of ADP at CII-CII interfaces opposite each other in the hexameric ring. KaiC protomers adjacent to these sites adopt a ‘compressed’ conformation that brings the CI and CII domains together (**Fig. 1C, 2A**), while the other four protomers remain in the extended conformation. It should be noted that C_2_-symmetry is rare in hexameric AAA+ proteins, with only a few other examples previously reported (*15-17*). In contrast to these previously described ‘dimer of trimers’ states, which each exhibit apo nucleotide pockets at their seam protomers (**Fig. S5)**, the C_2_-symmetric state of KaiC is unique in having its seam protomers occupied by ADP, suggesting thermodynamic coupling between CII phosphorylation and nucleotide state (see **Supplemental Text**). The ADP-bound protomers in our C_2_-symmetric KaiC structure also exhibit changes in accessibility of the pore loop (**Fig. S6A**), which is in turn linked to CII autophosphorylation (*18*) (**Fig. S6B-C**), providing a mechanistic link between KaiA association and nucleotide exchange in CII.

**Fig. 2.**
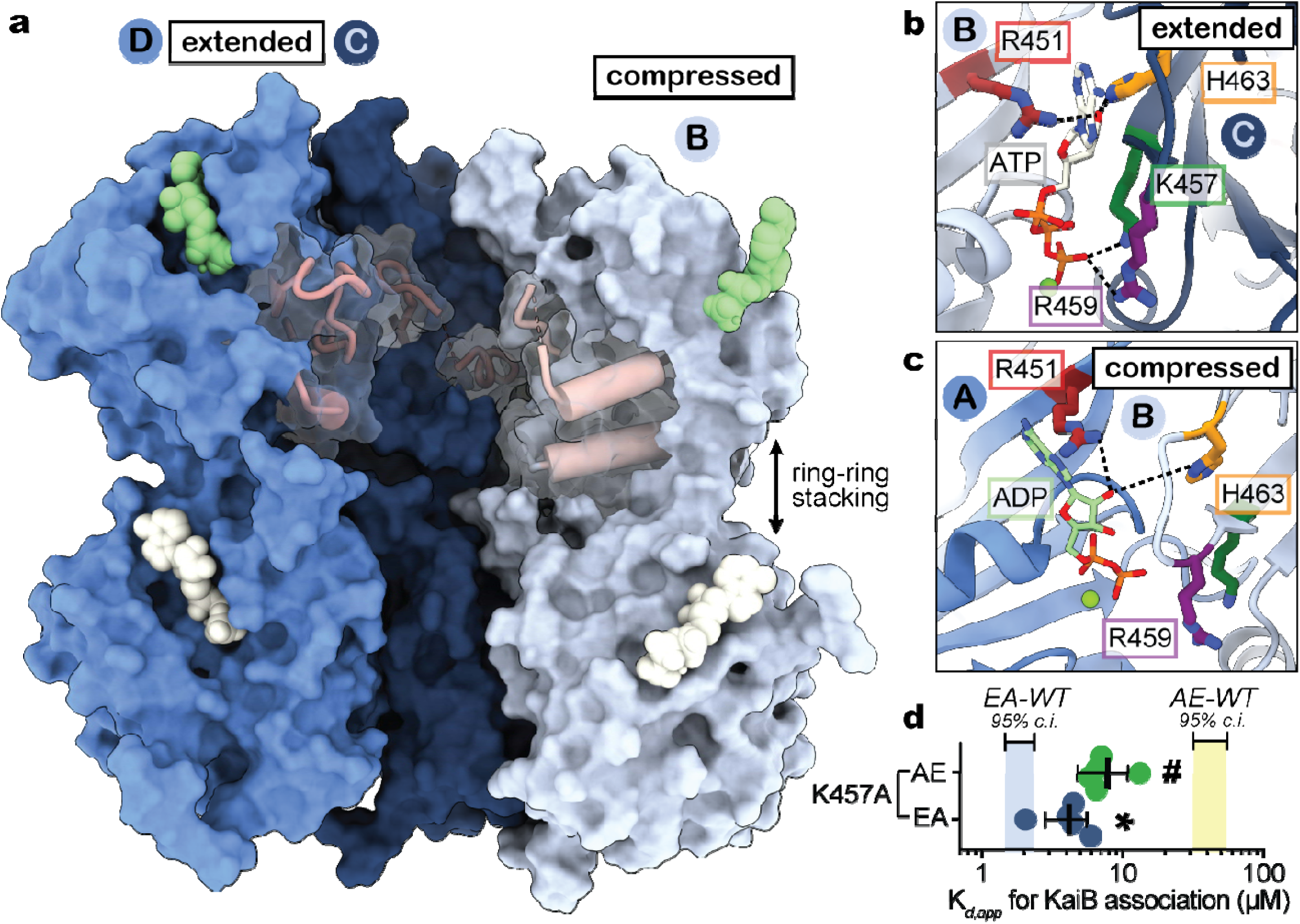
Nucleotide interactions at CII-CII interfaces govern the transition between expanded and compressed conformations. **a**) Surface representation of KaiC protomers in the C_2_-symmetric state with CII-α8 and CII-α9 depicted as pink cylinders. **b**) Closeups of the CII nucleotide interface in the expanded and (**c**) compressed KaiC protomer conformations. **d**) K_*d,app*_ for the interaction of KaiB with daytime (green) and nighttime (dark blue) KaiC variants bearing the K457A substitution (n = 5 with standard deviation). Symbols (**#**, P > 0.0001; *, P > 0.01) depict the results of unpaired parametric t-tests comparing each mutant to its respective unsubstituted phosphomimetic.

These observations raise the question: how does ring compression influence KaiB association? In the extended KaiC protomers of both the C_6_ and C_2_-symmetric structures, the sidechains of residues K457 and R459 interact directly with the *γ*-phosphate of ATP (**Fig. 2B**). However, at the ADP-occupied CII interfaces, contact with these sidechains is lost (**Fig. 2C**) and the CII domain on the compressed protomer breaks away from the adjacent CII interface. Computational studies have highlighted the importance of nucleotide interactions at CII interfaces on global conformational dynamics of KaiC (*19*). Given the apparent coupling between CII nucleotide interactions and the configuration of *cis* interacting domains in the compressed protomers, we wondered whether introducing an alanine residue in place of K457 would make this state accessible to the daytime variant and influence KaiB association. Consistent with our prediction, the K457A mutation resulted in a ∼6-fold increase in KaiB affinity for daytime KaiC (**Figs. 2D, S1B**), showing that KaiB association is bolstered by mutations that promote the compressed conformer in the daytime state. Likewise, a smaller effect was observed with the same mutant in the nighttime variant, demonstrating that K457A enhances KaiB association via dynamic interactions in the CII nucleotide cleft.

CII nucleotide state and associated ring compression are also linked with changes in secondary structure in the C_2_-symmetric KaiC structure. In particular, the α9-helix, where the adjacent phase-determining phosphosites S431 and T432 are located, comprises only a single turn in the extended protomers (**Fig. 3A**), but was lengthened significantly in the ADP-bound, compressed protomers (**Fig. 3B**). When we inhibited helical elongation by mutating I430 to glycine, we observed a ≥50-fold decrease in KaiB affinity (**Fig. 3C, S1C**), demonstrating that KaiB association is dependent on secondary structure formation induced by the proximal phosphosites.

**Fig. 3.**
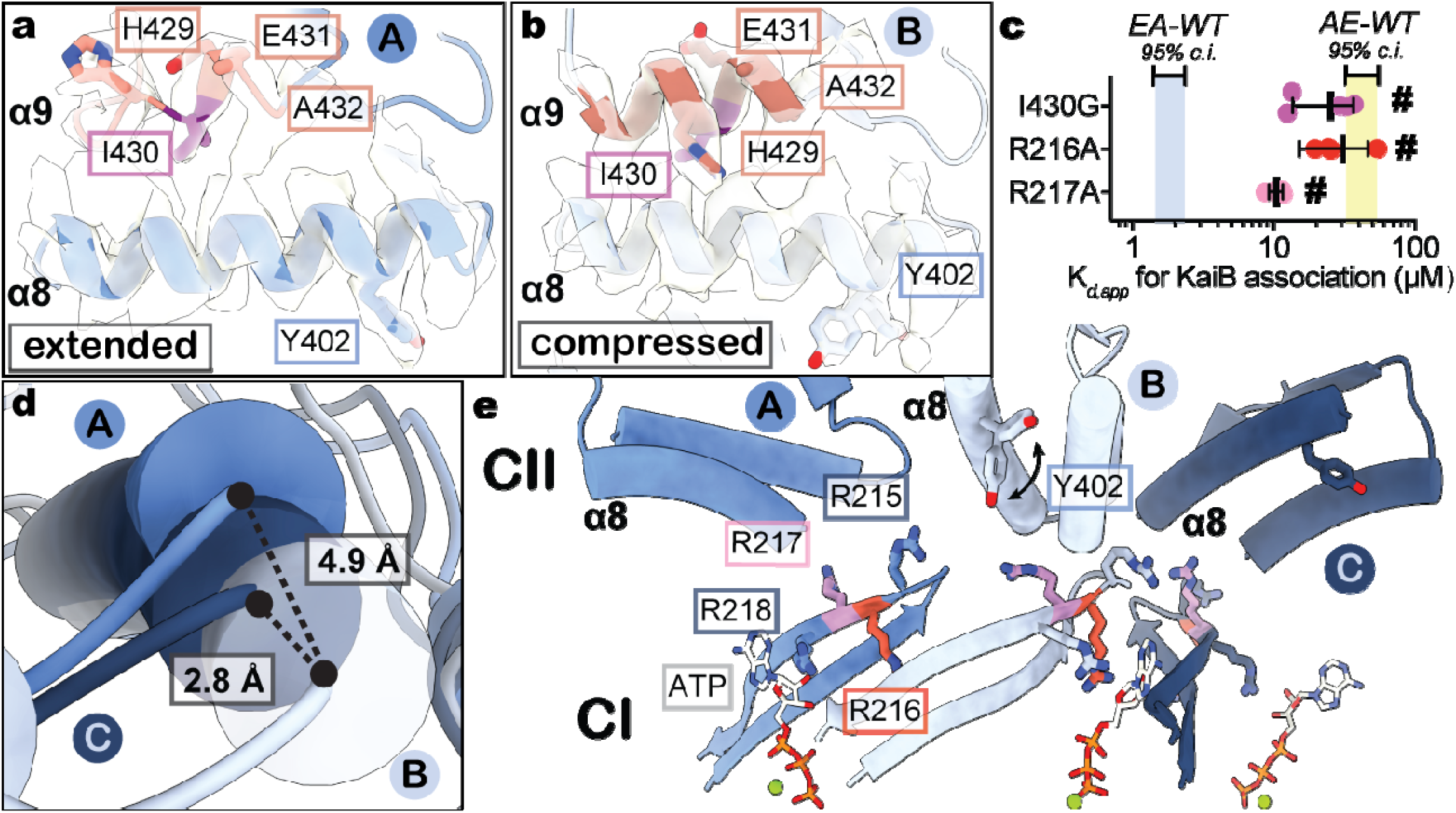
Compression of the *cis-*CI-CII interface primes nighttime KaiC for KaiB association. **a-b**) EM density and atomic models for residues S424 through T434 (pink) and CII-α8 helices in expanded (**a**, contour = 0.0343)) and compressed (**b**, contour = 0.0307) protomers of the C_2_-symmetric state of KaiC-EA. **c**) KaiB affinity for various mutants of CII-α9 and arginine tetrad (n ≥ 4 with standard deviation) in nighttime KaiC. Symbols represent one-way ANOVA (**#**, P > 0.0001) with Dunnett’s multiple comparisons for each mutant against KaiC-EA. **d**) Cylinder depictions of the CII-α8 helices from the three unique protomer conformations of the compressed hexamer, aligned about their CI domains. **e**) Proximity of CII-α8 to the arginine tetrad in the C_2_-symmetric state.

Lengthening of CII-α9 was also associated with translocation of CII-α8 toward the CI-CII interface in our compressed KaiC structure (**Fig. 3D**). A recent study highlighted the significance of tyrosine 402 as a key pacemaking residue on CII-α8 (*20*), with mutants at this position exhibiting extreme changes in CI ATPase activity and correlated circadian period *in vivo* (e.g., from 15 hours to >6 days). Notably, we observed EM density for distinct rotamer conformations of Y402 in the compressed protomers (**Fig. S7A**) situated near other phase-determining sidechains on the linker connecting CI and CII (*21*), indicating that dynamic interactions of CI with elements on CII-α8 are key determinants of circadian period for the oscillator.

The translocation of CII-α8 between the extended and compressed conformations (**Fig. 3D**) appears to regulate biochemical activity of CI via four sequential, highly conserved (**Fig. S8**), arginine residues in the CI domain known as the ‘arginine tetrad’ (**Fig. 3E**) (*10*). These residues are essential for circadian rhythms, forming a network of electrostatic interactions that join each CI protomer to both clockwise and counter-clockwise neighbors (*10, 22*) (**Fig. S9, Table S2**). The arginine tetrad is situated between the CI and CII domains on the tip of the CI-β9 hairpin that leads directly into the nucleotide binding pocket of CI (**Fig. S9B**), where enzymatic activity is required for KaiB association (*23*). To test whether the arrhythmic phenotype of arginine tetrad mutants (*10*) is due to disrupted regulation of KaiB binding, we measured KaiB affinity for two arginine tetrad mutants. Affinity was reduced by >10-fold for R216A (**Fig. 3C, S1C**) and by ∼6-fold for R217A, highlighting the importance of interdomain regulation for KaiB association.

The cooperative nature of KaiB binding to the KaiC hexamer is well established (*24, 25*). Given that the arginine tetrad is structurally poised to facilitate both *trans* and *cis* protomer regulation, we next wondered whether KaiC-R217A is capable of regulating cooperative recruitment of KaiB. To quantify KaiB cooperativity, we included unlabeled KaiB-I87A, often abbreviated as fold-switched or fsKaiB, as a secondary titrant to stimulate cooperative association of fluorescently labeled KaiB in KaiC binding assays (**Fig. S10A**) (*26*). Using this approach, we measured a cooperativity index of ∼25 ± 2 for KaiC-EA with fsKaiB (**Fig. 4A, S10B**). Since KaiB takes on the structure of fsKaiB when associated with KaiC (*8, 9, 27*), we interpret this as a ∼25-fold affinity enhancement for association of subsequent KaiB molecules once the first KaiB molecule binds to the hexamer. The R217A mutant showed a cooperativity index of only ∼1.6 ± 0.5, indicating that *trans* protomer communication traversing the arginine tetrad is required for KaiB cooperativity. Another arginine tetrad residue, R218, appears to be involved in *trans* regulation via interaction with the 2’ hydroxyl of the nucleotide at the CI-CI interface, sandwiched between the protomers by hydrogen bonds to R218 and H230 (**Figs. 4B, S11, Table S2**). Substitution of H230 with alanine also diminished cooperativity (**Fig. 4A**), despite having a relatively small (∼4-fold) effect on KaiB affinity (**Fig. 4C, S1D**), demonstrating that the CI active site is crucial for synchronizing KaiB recruitment on adjacent protomers.

**Fig. 4.**
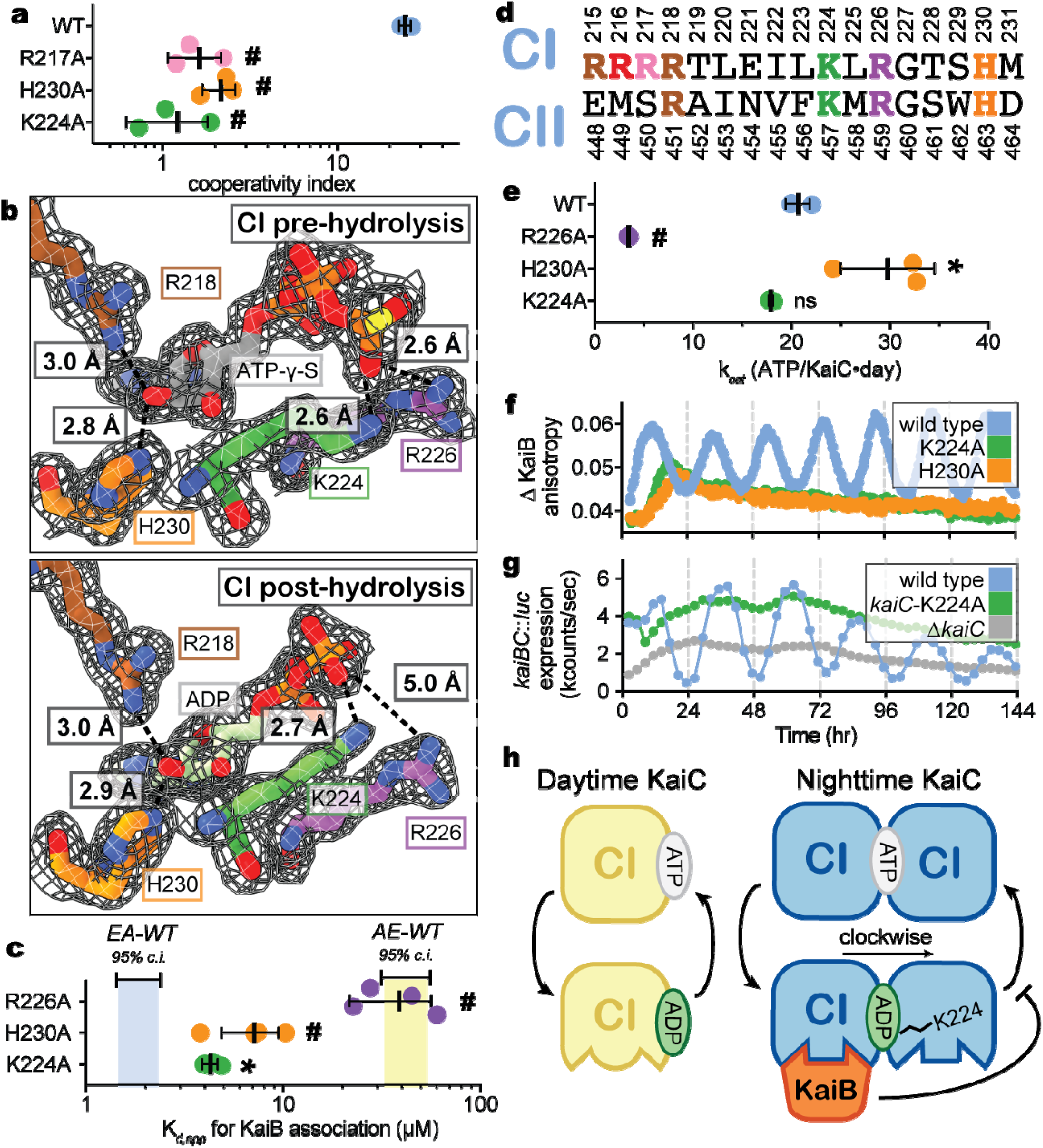
Interactions in the CI nucleotide binding pocket link ATP hydrolysis to cooperative KaiB recruitment and circadian rhythms. **a**) Cooperativity indices for KaiC-EA variants (n = 3 with standard deviation), determined with fsKaiB as secondary titrant (see text). Symbols represent one-way ANOVA (ns, not significant; *: P > 0.01; **#**, P > 0.0001) with Dunnett’s multiple comparisons to wild-type KaiC-EA. **b**) Atomic models and electron density (F_c_-F_0_, *σ* = 0.21) for nucleotide interactions from the pre-hydrolysis (PDB 4TL8, top) or post-hydrolysis (PDB 4TLA, chain C, bottom) CI domain crystal structures (*22*). **c**) *K*_*d,app*_ of KaiB binding to nighttime KaiC variants (n ≥ 4 with standard deviation) and statistical analysis as described for (**a**). **d**) Alignment of CI and CII protein sequences from *S. elongatus*. **e**) Turnover rate constants (*k*_*cat*_) for ATP hydrolysis by KaiC-EA variants (n = 3 with standard deviation) with statistical analysis as described for (**a**). **f**) *In vitro* oscillation reactions for KaiC mutants in the presence of KaiA and KaiB. **g**) Representative bioluminescence timecourses from a luciferase reporter gene driven by P*kaiBC* in *S. elongatus* cultures entrained under 12-h light/dark cycles for 48-hr and subsequently allowed to free-run in constant light (n = 6). **h**) Schematic model for the role of CI ATPase activity in regulation of KaiB association to KaiC.

It should be noted that CI residues R218 and H230 have analogs in CII, R451 and H463, that play similar structural roles in *trans* interactions in the CII domain via the nucleotide (**Fig. 2B**). Conversely, R215, R216 and R217 of the arginine tetrad are not conserved in CII (**Figs. 4D, S8**), indicative of their role in CI-CII interdomain regulation in *cis*. However, the overall architecture of the nucleotide binding pocket is strikingly well-conserved between the CI and CII domains of KaiC, and nearly invariant in KaiC across cyanobacteria (**Fig. S8**). In particular, the CI and CII domains both contain a lysine/arginine pair (K224 and R226 in CI, K457 and R459 in CII). Previously reported high-resolution structures show that K224 changes orientation to coordinate the ultimate nucleotide phosphate at both pre- and post-hydrolysis CI-CI interfaces (**Fig. 4B, S11, Table S3**), whereas R226 coordinates nucleotide only in the ATP-bound state (*22*). To determine whether these interactions are required for ATP hydrolysis, we measured the enzymatic turnover rate, *k*_*cat*_, of the R226A, K224A, and H230A mutants (**Figs. 4D, S12**). We found that the R226A mutation drastically decreased both ATPase activity and KaiB affinity, similar to other catalytically dead variants (*23*). Conversely, the H230A mutation increased ATPase activity relative to wild type, but diminished KaiB affinity (**Fig. 4E**), suggesting a more nuanced coupling between ATPase activity and KaiB association than simple correlation. Consistent with this, the K224A mutant had the smallest effect on KaiB affinity and ATPase activity of all the CI active-site mutants tested, but it essentially eliminated cooperativity (**Fig. 4A**), indicating that K224 is dispensable for KaiB association on its own protomer, but critical for cooperative regulation of KaiB recruitment in *cis*.

Given its apparent structural role in sensing nucleotide state at the clockwise CI-CI interface, this result suggests that K224 plays a key role in coupling ATPase activity to cooperativity in KaiB association, thus imparting a switch-like nature to the assembly of protein complexes––a function that has been well documented for other RecA-like AAA+ proteins (*28*). Notably, we observed total loss of circadian rhythms *in vitro* with KaiC variants lacking cooperativity (**Figs. 4A, 4F**), demonstrating that the core oscillator cannot function in the absence of cooperative KaiB recruitment. Furthermore, the KaiC-K224A variant did not support circadian gene expression in *S. elongatus* (**Fig. 4G**), demonstrating that the presence of additional factors cannot compensate for the loss of CI nucleotide sensing and *cis* regulation of KaiB recruitment in *vivo*.

It should be noted that although ATP hydrolysis in CI is required for KaiB association (*23*) and ADP-bound CI interfaces are observed in KaiB-bound KaiC structures (*8, 9*), we never observed ADP-bound CI domains in our cryo-EM structures, indicating that phospho-dependent regulation of KaiB association by CII does not occur simply by trapping the ADP-bound CI state prior to KaiB association. Rather, we propose that ADP is replaced with ATP after hydrolysis unless KaiB association locks the CI domain into the post-hydrolysis state by inhibiting ADP release (**Fig. 4H**). In this model, S431 phosphorylation regulates the CI domain through elongation of the CII-α9 helix, leading to *cis* domain compression and translocation of CII-α8 axially towards the arginine tetrad on CI. There, dynamic interactions regulate KaiB association by coupling CI nucleotide state with KaiB occupancy on the same and adjacent CI domains.

This work provides a clear mechanistic link between the daily phosphorylation cycle of KaiC and quaternary structural changes that mediate negative feedback in the cyanobacterial circadian pacemaker. Although CI ATPase activity is correlated with circadian period *in vivo* and *in vitro* for many KaiC mutants (*20-22*), no precise chemo-mechanical mechanism has yet been described to explain this. Our results demonstrate that phospho-dependent coupling between the ATPase cycle on CI and *cis* domain compression influences clock assembly through cooperative KaiB recruitment, explaining why the most extreme period-altering mutations in KaiC are located on CII-α8 or nearby on the linker that connects the CI and CII domains (*20, 21*). These structures and the associated biochemistry pave the way for a comprehensive understanding of the post-translational oscillator from both modeling and reverse-engineering perspectives, and underscore the elegant complexity of this sophisticated and primeval timepiece.

## Acknowledgments

We thank J.C. Ducom at Scripps Research High Performance Computing for computational support, as well as B. Anderson at the Scripps Research Electron Microscopy Facility for microscopy support.

## Author contributions

Conceptualization: JAS, CRS, GL, CLP

Investigation: JAS, CRS, AGC, AMF, DE, CS, DCE, JGP

Funding acquisition: SSG, AL, GL, CLP

Project administration: JAS, GL, CLP

Supervision: SSG, AL, GL, CLP

Writing––original draft: JAS, CRS, GL, CLP

Writing––review & editing: JAS, CRS, AMF, SSG, AL, GL, CLP

## Data and materials availability

KaiC structures have been deposited in the PDB and EMDB with the following codes: EA Compressed State, PDB 7S65 and EMD-24850; EA Expanded State, PDB 7S66 and EMD-24851; AE Expanded State, PDB 7S67 and EMD-24852. All data are available in the main text or the supplementary materials

## Supplementary Materials

Materials and Methods

Supplementary Text

Figs. S1 to S12

Tables S1 to S6

References (*29*–*55*)

Data S1 to S3

