## Supplementary Material for "Hidden conformations differentiate day and night in a circadian pacemaker"

**This PDF file includes:**

Materials and Methods
Supplementary Text
Figs. S1 to S12
Tables S1 to S6
Captions for Data S1 to S3
Supplemental References 1-55

**Other Supplementary Materials for this manuscript include the following:**

Data S1 to S3

### Materials and Methods

#### Site-directed mutagenesis, protein expression and purification

All protein constructs were expressed from the Kanamycin-resistant pET-28b plasmid, with an N-terminal His-SUMO tag for affinity purification. Mutations were introduced by long-range amplification of the whole vector, installing point mutations via oligonucleotide primers in the method originally described by Liu et al. (*29*). KaiA and KaiB were prepared as described previously (*7, 13*). KaiC constructs were expressed in BL21 (DE3) *E. coli* cells (NEB). Cultures were grown in M9 medium under antibiotic selection. Cells were grown to an optical density ($\lambda$ = 600 nm) between 0.4 and 0.8 by shaking at 37°C, and were subsequently cooled to 18°C before induction with 200 µM IPTG and overnight shaking at 18°C.

Next, the cultures were spun down and resuspended in 50 mM $NaH_2PO_4$ pH 8.0, 500 mM NaCl, 20 mM Imidazole, 5 mM β-mercaptoethanol, 1 mM ATP and 1 mM $MgCl_2$ ($Ni^{2+}$ Buffer A). For KaiC, all buffers also contained 1 mM ATP. Cell suspensions were subsequently either frozen for later purification, or lysed immediately by passing through an Emulsiflex high pressure homogenizer (Avestin) ≥ 10 times at ≥ 10,000 psi. The soluble portion of the lysate was then recovered by centrifugation for 45 minutes at 45,000 rcf at 4°C. The clarified lysate was loaded onto Ni-NTA agarose (Qiagen, 5 mL per 2 L of initial E. coli culture) that had been equilibrated with 10 column volumes of $Ni^{2+}$ Buffer A.

The Ni-NTA resin loaded with lysate was washed with 10 column volumes of $Ni^{2+}$ Buffer A and bound proteins were eluted with 30 mL (per 5 mL Ni-NTA) of $Ni^{2+}$ Buffer containing 250 mM Imidazole. The eluted protein was then treated with ~10 nmol Ulp1 enzyme for ≥ 30 minutes at room temperature to cleave the SUMO tag. Cleaved KaiC was then concentrated to ≤ 2 mL using a 30 kDa centrifugal filter (Millipore) and injected to an Superdex 200 gel filtration column (GE Healthcare) equilibrated with 20 mM Tris pH 7.4, 150 mM NaCl, 1 mM ATP, 1 mM $MgCl_2$, and 1 mM TCEP (KaiC buffer). Fractions containing KaiC hexamer were then quantified using absorbance readings, frozen in liquid nitrogen, and stored at -70°C.

#### KaiB binding assays

Binding assays were performed, generally, as described in (*26*). Briefly, 30 µL of KaiC buffer containing 0.1% Tween and 50 nM fluorescently labeled KaiB was pipetted into 21 adjacent wells of a 384-well flat-bottom black polystyrene assay plate (Corning). KaiC aliquots were then thawed and supplemented with 0.1% Tween and 50 nM fluorescently labeled KaiB before 90 µL portions were added to the plate in line with the 21 buffer wells. Serial dilution was then executed by transferring 60 µL of KaiC mixture to the first buffer well, mixing by pipetting in and out ≥ 6 times and then transferring 60 µL from that well to the subsequent well and repeating this process for all 21 wells.

The plates were then sealed and incubated overnight (~15 h) at room temperature. The plates were then uncovered and fluorescence anisotropy was measured using a SYNERGY2 microplate reader (BioTek). 10-20 measurements were collected and averaged for each well, and data were analyzed using Prism (GraphPad). Data were fit to the Langmuir isotherm:

$$a = \Delta a_{total} * \frac{[KaiC]}{[KaiC] + K_{d,app}} + BG$$

Where a is anisotropy, Δa is the change in anisotropy over the course of the experiment, [KaiC] is the concentration of KaiC variant, $K_{d,\mathrm{app}}$ is the apparent dissociation equilibrium constant and BG is the background anisotropy. In cases where saturation could not be reached, as evidenced by a stable anisotropy value as a function of KaiC, a lower limit for $K_{d,app}$ was estimated by holding $\Delta a_{\mathrm{total}}$ constant at 0.043 anisotropy units.

The raw anisotropy values were subsequently converted to change in anisotropy by subtracting the *BG* term from the anisotropy values in each titration curve (see **Data S1**). The reported $K_{d,app}$ values were then determined by refitting the background subtracted Δ anisotropy values to the Langmuir function above with the *BG* term omitted.

Electron microscopy

*Sample Preparation*

KaiC-EA or KaiC-AE was briefly incubated on ice in 20 mM Tris pH 7.4, 150 mM NaCl and 1 mM each of ATP, $MgCl_2$ and TCEP. A sample (2.5 µL at 1.5 mg/ml) was then applied to an UltraAuFoil R1.2/1.3 300-mesh grid (Electron Microscopy Services), which was previously plasma-cleaned using a Gatan Solarus (75% argon/2% oxygen atmosphere, 15 W for 7 seconds). To overcome an orientation bias in ice, an additional dataset was obtained by applying KaiC-EA (2.5 µL at 0.2 mg/ml) to holey C-flat grids, which previously had thin carbon floated on them. These grids were pretreated with 5 µL of 0.1 % (w/v) poly-L-lysine hydrobromine (Polysciences), blotted to dryness and then washed 3X with 10 µL of water. Grids with applied sample were then manually blotted with filter paper (Whatman No.1) for ~3 seconds in a 4°C cold room before plunge-freezing in liquid ethane cooled by liquid nitrogen.

For the detergent dataset, KaiC-AE was incubated on ice with 4 mM fluorinated fos-choline-8 (Anatrace) added to the sample buffer. 2.5 µL of 6 mg/ml sample was added to UltraAuFoil R1.2/1.3 300-mesh grids (Electron Microscopy Services) that were pretreated with the addition of a graphene monolayer, using a modified protocol (*30*) for the deposition of graphene and made hydrophilic via UV/ozone cleaner (UVOCS T10 x 10 system) as previously described (6), and then incubated for 4 minutes in a 4°C cold room to concentrate particles on the surface of the grid before manual blotting for ~3 seconds followed by plunge freezing into liquid ethane.

*Data acquisition*

All cryo-EM data were acquired using the Leginon automated acquisition software (*31*). For KaiC-EA, all real-time image pre-processing, consisting of frame alignment, local CTF estimation, and particle picking were performed using the Appion image-processing pipeline during data collection (*32*). For KaiC-AE, all preprocessing was performed using Relion 3.0 (*33*)

Image collection was conducted using a Thermo Fischer Talos Arctica operating at 200 KeV and equipped with a Gatan K2 Summit DED. We used a nominal magnification of 36,000X, corresponding to a pixel size of 1.15 Å at the detector. For KaiC-EA, 1,137 movies were collected at a 40° tilt to overcome preferred orientation (*34*) with an exposure time of 8 seconds and an exposure rate of 7.94 $e^-$/pixels/s and a total exposure of 48 $e^-$/Å$^2$ (0.6 $e^-$ per frame) with a nominal defocus range of 0.8 – 1.7 µM. For the thin carbon KaiC-EA dataset, we collected 8,405 movies with no tilt and with an exposure time of 9 seconds. We used an exposure rate of 7.5 $e^-$/pixels/s for a total exposure of 51 $e^-$/Å$^2$ (1.4 $e^-$ per frame) with a nominal defocus range of 0.5 – 1.5 µM. In the tilted KaiC-AE dataset, 1,132 movies were collected at 40° tilt with an exposure time of 11 seconds and exposure rate of 5.71 $e^-$/pixels/s for a total exposure of 48 $e^-$/Å$^2$ (0.9 $e^-$ per frame) and with a nominal defocus range of 0.8 – 1.7 µM.

For the KaiC-AE detergent dataset, 1,541 movies were collected with no tilt, using an exposure time of 6.2 seconds and exposure rate of 8.61 $e^-$/pixels/s for a total exposure of 40.32 $e^-$/Å$^2$ (0.7 $e^-$ per frame) with a nominal defocus range of 0.5 – 2.0 µM.

*Image processing*

Movies of all KaiC datasets were aligned and dose-weighted in the Appion pipeline using Motioncorr2. For the tilted KaiC-EA dataset, 239 images were used for automated particle picking using Difference of Gaussians (DoG) picker to yield an initial 151,640 particles (Fig. 5.3a) (*35*). Gctf was used for local CTF estimation, using an increased raster spacing of 500 to accommodate the defocus gradient resulting from the tilted data collection (*36*). Fourier-binned 4x4 particles were then subjected to reference-free 2D classification with multivariate statistical analysis (MSA) and multi-reference alignment (MRA) in the Appion pipeline (*32*). The two best classes representing tilted and side views were selected for template-based particle picking using FindEM (*37*), resulting in 2,016,826 particle picks from the entire dataset.

For initial particle curation, the ~2 million particle picks were subjected to two rounds of reference-free 2D classification in Relion (*33*), leaving 1,341,217 particles bearing structural details. A previous crystal structure of KaiC (pdb: 3K0C) was used to generate an initial model using the molmap feature in UCSF Chimera (*38*). This model was low-pass filtered to 20 Å resolution for 3D autorefinement in Relion, which resulted in a ~9.6 Å map that did not have discernable density for the majority of the CII ring. 3D classification into 6 classes without alignment resulted in two classes with mostly intact CII and CI rings, totaling 579,706 particles. These particles were re-centered and re-extracted at 2.3 Å/pixel.

For the thin carbon dataset, FindEM was used along with the same tilted and side view templates from the titled dataset, resulting in 2,856,972 particle picks. CTFFIND4 was used for CTF estimation prior to extracting particles. Particles were Fourier-binned 4×4 and then subjected to one round of 2D classification in Relion, allowing for the removal of false particle picks and for the identification and selection of side views. 2,856,972 particles that belonged to exemplary side views were re-centered and re-extracted at 2.3 Å/pixel and then combined with the 579,706 particles from the tilted dataset. 3D classification in Relion with alignment, asking for four

classes, gave a single well-resolved class comprising 1,002,777 particles. 3D autorefinement of the well-resolved class resulted in a map with a global resolution of ~4.7 Å. These particles were re-centered and re-extracted at 1.15 Å/pixel before undergoing a second 3D autorefinement, giving a new map with a global resolution of ~4.1 Å. Additional 3D classification, without alignment and using a Tau value of 10 and asking for four classes, generating three high resolution classes – two seemingly $C_2$-symmetric and one seemingly $C_6$-symmetric.
The 658,212 particles belonging to the two seemingly $C_2$-symmetric classes were joined for one round of 3D classification into three classes with no alignment and a Tau value of 20. One class containing the highest resolution features was selected for further processing. These 122,855 particles underwent 3D autorefinement with applied $C_2$ symmetry, giving a map with a global resolution of ~3.7 Å. An in-house python script was used to group particles by image shift, which then underwent iterative rounds of CTF refinement in Relion 3.1 refining both beam tilt and per-particle astigmatism correction. Subsequent 3D autorefinement with applied $C_2$ symmetry and a soft mask around the CI ring resulted in a map at ~3.5 Å. Postprocessing in Relion yielded a final $C_2$-state map at ~3.2 Å global resolution.

Particles from the seemingly $C_6$-symmetric class from the earlier 3D classification without alignment underwent 3D autorefinement, resulting in a ~4.1 Å map with 91,869 particles. 3D autorefinement with applied $C_6$ symmetry returned a ~3.7 Å map. Image shift grouping and iterative CTF refinement in Relion 3.1, followed by 3D autorefinement with applied $C_6$ symmetry and a soft mask around the CI ring resulted in a ~3.2 Å map. Postprocessing resulted in a final map for the $C_6$-state at ~2.8 Å resolution.

Particle picking for the detergent dataset for KaiC-AE was done using FindEM template picking, using a single axial and side view from the earlier datasets as templates, resulting in 427,620 particle picks. Particles were extracted at 2.3 Å/pixel and then underwent 2D alignment with cryoSPARC. Particles belonging to classes that showed secondary structure were selected as input particles for *ab initio* reconstruction, asking for 3 classes. One class, containing 111,877 particles, resembled the stacked ring shape of KaiC, and was selected for homogeneous refinement resulting in a map at ~6.3 Å resolution. These particles then underwent a second round of ab initio reconstruction with 3 classes, with one class containing 57,067 particles that resolved to ~6.7 Å after homogenous refinement. These particles were subjected to one final round of ab initio reconstruction with 2 classes, resulting in one class showing detailed structural features, consisting of 42,838 particles. Homogeneous refinement using $C_1$ symmetry resulted in a ~6.5 Å reconstruction, followed by homogenous refinement with $C_6$ symmetry, yielding a map at ~4.8 Å resolution. The full dataset (427,620 particle picks) was extracted, unbinned at 1.15 Å/pixel, and underwent 2D classification in Relion. The selected 367,247 particles were then subjected to 3D classification in Relion with a limited resolution E-step of 7 Å, and using the ~4.8 Å map obtained from cryoSPARC as an initial model. Of the 4 resulting classes, one was selected for further processing with 89,892 particles. These particles underwent 3D autorefinement, resulting in a ~5.6 Å map, followed by CTF Refinement and 3D autorefinement once again, resulting in a ~4.8 Å map. Another round of CTF Refinement was conducted, followed by 3D autorefinement with $C_6$ symmetry resulting in a ~4.2 Å map which led to a final sharpened map at ~3.8 Å after postprocessing. All reported resolution are according to the FSC at a cutoff of 0.143.

*Atomic model building and refinement*

For both the expanded and compressed state models of KaiC-EA, a crystal structure of the phosphomimetic mutant S431A/T432E was used as a starting point for model building (PDB: 3K0C, (*39*)) with the C-terminal A-loops deleted. For the $C_2$-state, which showed large-scale repositioning of the C-terminal domains, proSMART local restraints (*40*) were generated in Coot (*41*) which were then used to refit the domains into the cryo-EM density. After one round of real-space refinement in Phenix using default parameters and 5 macrocycles (*42*), Coot and ISOLDE (*43*) was used to improve main chains and side chains. The Molprobity server (*44*) (http://molprobity.biochem.duke.edu/) and PDB validation service server (https://validate-rcsb-1.wwpdb.org/) were used to identify problem regions for subsequent correction in Coot.

KaiC autophosphorylation assays

Samples (1 mL) of WT or mutant KaiC proteins were prepared in KaiC buffer. Samples of KaiC were mixed with KaiA to generate a final concentration of 1.5 μM KaiA and 3.4 μM KaiC. Samples were incubated at 30°C for 24 h, at which point 50 μL was removed and quenched with 10 μL of 6X SDS buffer and then incubated at 95°C for 5 minutes.

Phos-Tag™ acrylamide gels were prepared with a resolving gel consisting of 10% (w/v) acrylamide (29:1 acrylamide/bis-acrylamide) containing Tris-HCl pH 8.8 and supplemented with 50 μM Phos-tag™ reagent and 100μM $Mn^{2+}$. Gels were run at constant 25 mA with a 165 V limit for 35 minutes before loading any samples into the gel. Samples (10 μL) were loaded and run with the same mA and voltage parameters stated above for 170-190 minutes. Gels were stained using SYPRO® Orange following the Bio-Rad protocol. Band density was analyzed using ImageJ software (*45*). Error was estimated as the standard deviation from n = 3 replicate measurements analyzed separately.

KaiC ATPase assays

After the initial protein purification, frozen aliquots of each KaiC variant were thawed and run over a Superdex 200 gel filtration column (GE Healthcare) to exchange them into fresh buffer containing 50 mM MOPS pH 7.4, 150 mM $NaCl_2$, 1 mM TCEP, 1 mM $MgCl_2$, 400 μM ATP (ATPase buffer). Fractions containing KaiC hexamer were diluted to working concentrations with ATPase buffer for a final volume of 50 μL. Samples were then incubated at 30°C and 5 μL aliquots were removed at 0, 4, and 8-h timepoints and quenched using the ADP-Glo assay kit (Promega) according to manufacturer's instructions. Luminescence measurements (RLU values) were taken at room temperature with a SYNERGY2 microplate reader (BioTek) in opaque black 384-well microplates. Data analysis was performed using Prism (GraphPad).

Standard curves were generated each day in ATPase buffer containing 20%, 40%, 60%, 80%, and 100% ADP while maintaining 400 μM total nucleotide concentration with ATP. Aliquots (5 μL) of these samples were mixed with ADP-Glo reagent as described above. Plotting RLU versus μM ADP and solving for the x-intercept gives an equation which converts RLU values to μM ADP.

Because the concentration of substrate (ATP) was present in vast excess of the $K_M$ of 2 μM (*46*), the Michaelis-Menten equation simplifies to the following:

$$v_0 = k_{\text{cat}} \times [enzyme]$$

and the slope of the $v_0$ is the initial velocity, $k_{\text{cat}}$ is the catalytic turnover rate and [enzyme] is, in this case, the concentration of KaiC variant in the assay. Thus, we plotted $v_0$ for each concentration of a given mutant against [KaiC] and, using the equation above, extracted $k_{\text{cat}}$ as the slope of the resulting line.

Thermodynamic modeling of cooperativity indices

KaiC titrations were performed as described above, but with various concentrations of KaiB-I87A (fsKaiB) in both the KaiC and diluent stocks. Background anisotropy was subtracted from these data before analysis. Least squares fitting was performed using DynaFit (BioKin) as described in (*26*), with modifications (see **Data S2** for DynaFit scripts).

Specifically, the assumption that $K_5 = K_1$, was replaced by the assumption that $K_5 = K_3$. The latter assumption was not appropriate in (*26*) because the KaiB-related protein SasA was used as a secondary titrant. Because wild-type KaiB assumes the fold-switched conformation when in complex with KaiC, $K_5 = K_3$ is an appropriate assumption.

Anisotropy values were allowed to float initially, and then restricted for extraction of the final model. The two highest fsKaiB concentrations were also allowed to float when fitting data for the mutant KaiC variants. In some cases, particularly with KaiC-EA-WT, convergence could not be achieved for the absolute values of $K_2$ and $K_4$ (**Data S3**), though the ratio $K_2/K_4$ (which is equal to the reported cooperativity index for each variant $K_1/K_2$) was in good agreement between replicates.

*In vit*ro oscillation assays

KaiABC *in vitro* oscillator reactions were performed as described previously (*26, 47*). Briefly, purified protein stocks were mixed in buffer containing 20 mM Tris pH 8.0, 150 mM NaCl, 5 mM $MgCl_2$, and 1 mM ATP to a final concentration of 3.5 μM KaiC, 3.5 μM KaiB, 1.2 μM KaiA and 50 nM KaiB-K25C-fluorescein. Fluorescence anisotropy was monitored at 30°C using a CLARIOstar microplate reader (BMG Labtech). All data collection was performed using the fluorescein channel ($\lambda_{ex}$, 490 ± 5 nm; $\lambda_{em}$, 520 ± 5 nm) with a measurement taken every 15 minutes.

Strains and culture conditions for *in vivo* assays

Markerless incorporation of the K224A point mutation into *kaiC* of *Synechococcus elongatus* PCC 7942 was performed by CRISPR/Cas12a engineering as described previously (*48*). The plasmids and primers used for generating the kaiC-K224A strain are listed in **Table S4**.

Characterization of circadian rhythms in the *kaiC*-K224A strain were performed as described in (*26*). Briefly, bioluminescence was monitored from the P*kaiBC*::*luc* fusion reporter, inserted into a neutral site of the *S. elongatus* chromosome as previously described (*49*). Strains to be monitored were diluted to $OD_{750}$ = 0.2. 20 µL aliquots of the suspension were added to 280 µL pads of BG-11 agar with 3.5 mM firefly luciferin in 96-well plates. Cells were entrained under 12-h light-dark cycles (80 µmol $m^{-2}s^{-1}$) to synchronize clock phases. After 48 h of entrainment, cells were released into continuous light (30 µmol $m^{-2}s^{-1}$). Plates were transferred to a lighted stacker (40 µE light) attached to a Tecan Infinite M200 Pro and bioluminescence monitored every 2-3 h. All strains used in this study are listed in **Table S5**.

### Supplemental Text

Implications of $C_2$ symmetry on the mechanism of KaiC autophosphorylation and dephosphorylation

KaiC autophosphorylation is governed by interactions of the clock protein KaiA with C-terminal loops on KaiC, known as the A-loops, that bind to the central pore of KaiC when KaiA is absent (*18*). One characteristic feature of the $C_2$-symmetric state is the loss of A-loop interactions within the central pore (**Fig. S6A**). In the expanded state, the sidechain of E444 forms a hydrogen bond with the mainchain nitrogen of I490, located in the A-loop, in *trans*. However, this interaction is broken in the compressed conformation. Consistent with this, we observed constitutive autophosphorylation in the E444S pore-mutant, similar to prior studies where the A-loops were deleted entirely (*18*). Another consequence of the compressed state that occurs with $C_2$ symmetry is that A-loop availability is coupled between protomers opposite each other in the KaiC hexamer. This suggests coupling in the effect of KaiA on non-adjacent KaiC protomers, and helps to explain why sub-stoichiometric levels of KaiA are sufficient with respect to KaiC to maintain robust oscillation (*50*).

Furthermore, the thermodynamic coupling we observed across the KaiC hexamer extends to the CII nucleotide state, which has implications for the mechanism of autodephosphorylation, which proceeds via transfer of the phosphate from Serine 431 or Threonine 432 to the β-phosphate of ADP (*51*). As a result, CII autophosphorylation should be influenced by the availability of ADP in the CII active site via Le Chatelier's principle, and KaiA acts as a nucleotide exchange factor to exploit this characteristic (*52*). Thus, the observed structural link between nucleotide state and A-loop conformation suggests a direct coupling between KaiA association at the A-loops and accumulation of ADP in the CII active site.

It should be noted that regulation of phosphosites based on nucleotide state at CII is not unexpected given the sequence similarity between CI and CII (**Fig. S8**) and the fact that the conformational changes associated with the ADP-bound site in CI are centered on a homologous α-helical region (*22*) containing a negatively charged residue in place of S431.

Analogous roles of KaiC K224 to active site elements in other AAA+ proteins

The apparent role of KaiC K224 in coupling of ATP hydrolysis to a switch-like biochemical activity bears a striking similarity to a recently reported 'arginine coupler' in the bacterial helicase loader DnaC (*53*). In this system, mutation of a positively charged amino acid (the 'arginine coupler') just upstream of the arginine finger to alanine increased ATPase activity while eliminating the switch-like loading activity of DnaC. This motif is also conserved in mammalian initiator ATPases, and our observation of a similar mechanism in the cyanobacterial clock protein KaiC suggests that linking ATP hydrolysis to a state-switching activity could be more widespread in the AAA+ family.

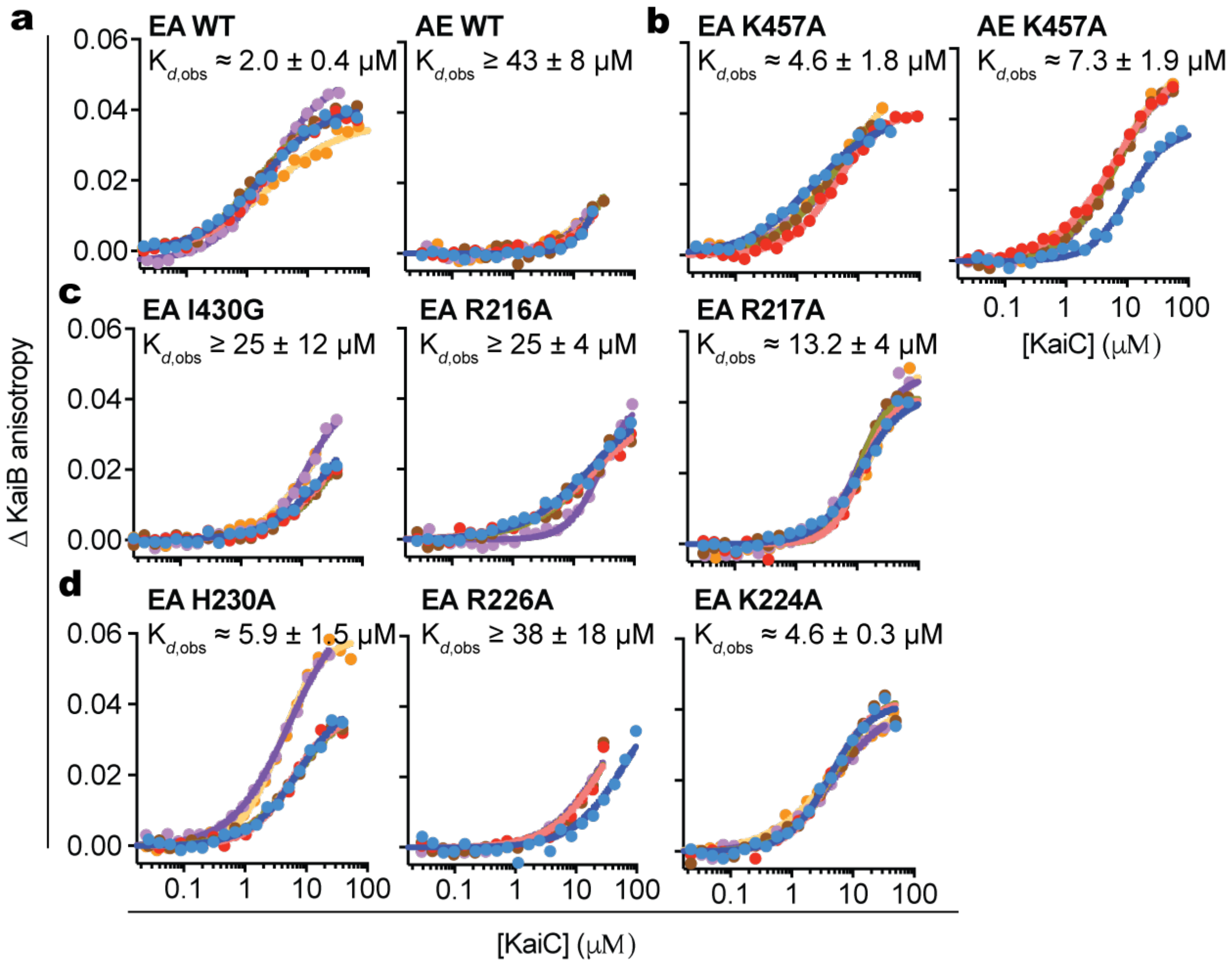


**Fig. S1: Affinity measurement for KaiC variants**. **a**) Equilibrium titrations of fluorescently labeled KaiB with KaiC variants and associated binding curves determined as described in Methods. Mean $K_{d,obs}$ values and standard deviation for n ≥ 4 replicate measurements are shown for each variant. Various colors are used to distinguish the individual replicates and binding curves from each individual experiment. Separate panels distinguish the data found in Figure 1, **b**) Figure 2, **c**) Figure 3 and **d**) Figure 4.

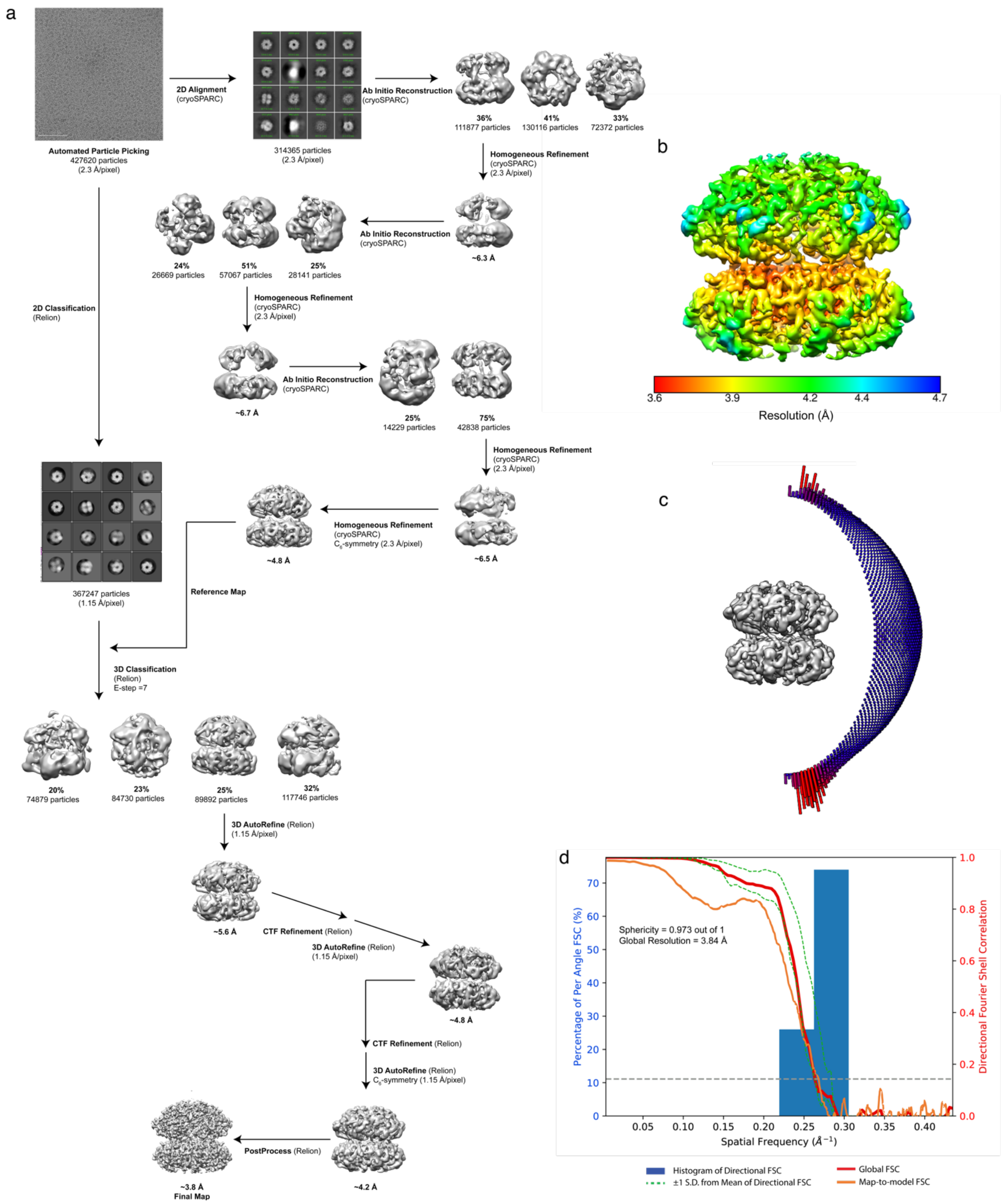


**Fig. S2. Image processing pipeline and validation of $C_6$ symmetric KaiC-AE structure. a**) Image processing pipeline for KaiC-EA datasets with fluorinated fos-choline-8. **b**) Local resolution estimation of cryo-EM reconstruction calculated by RELION (*33*). **c**) Euler distribution plots depicting particle orientations present in final reconstruction. More populated views are colored in red while less populated are depicted in blue. **d**) 3D Fourier Shell Correlation (3DFSC) (*34*) of final daytime state reconstruction, with a global resolution of 3.8 Å at FSC=0.143.

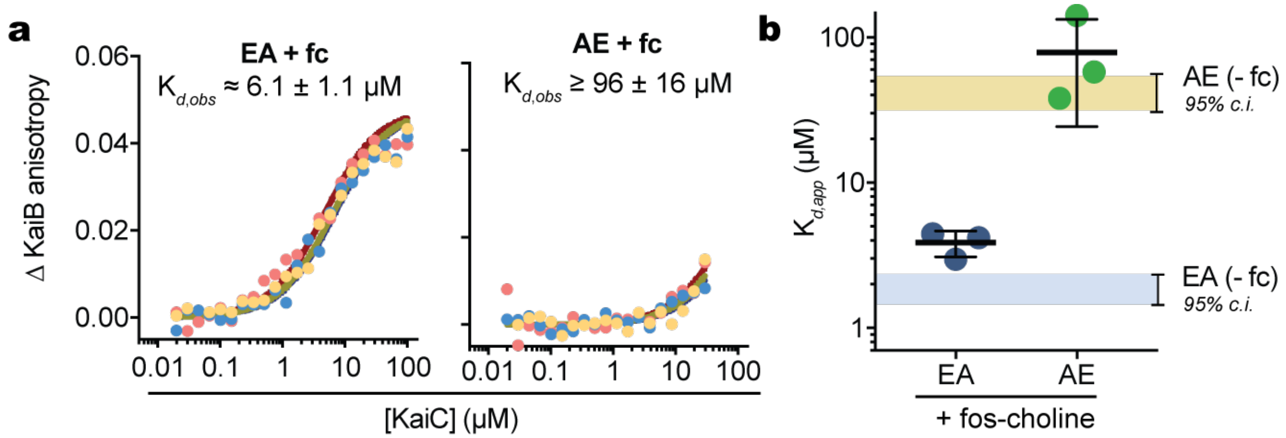


**Figure. S3. Discrimination of KaiC phosphomimetics for KaiB association persists in the presence of fluorinated fos-choline-8.** **a**) Equilibrium binding assays of nighttime and daytime KaiC in the presence of 4 mM fluorinated fos-choline-8. Triplicate measurements are shown in red, yellow and blue with binding curves determined as described in methods. **b**) Apparent $K_{DS}$ from binding assays with (circles with mean and error bars representing standard deviation) and without (horizontal bars representing the 95% confidence interval of the $K_{d,obs}$ from the main manuscript) the presence of fluorinated fos-choline-8.

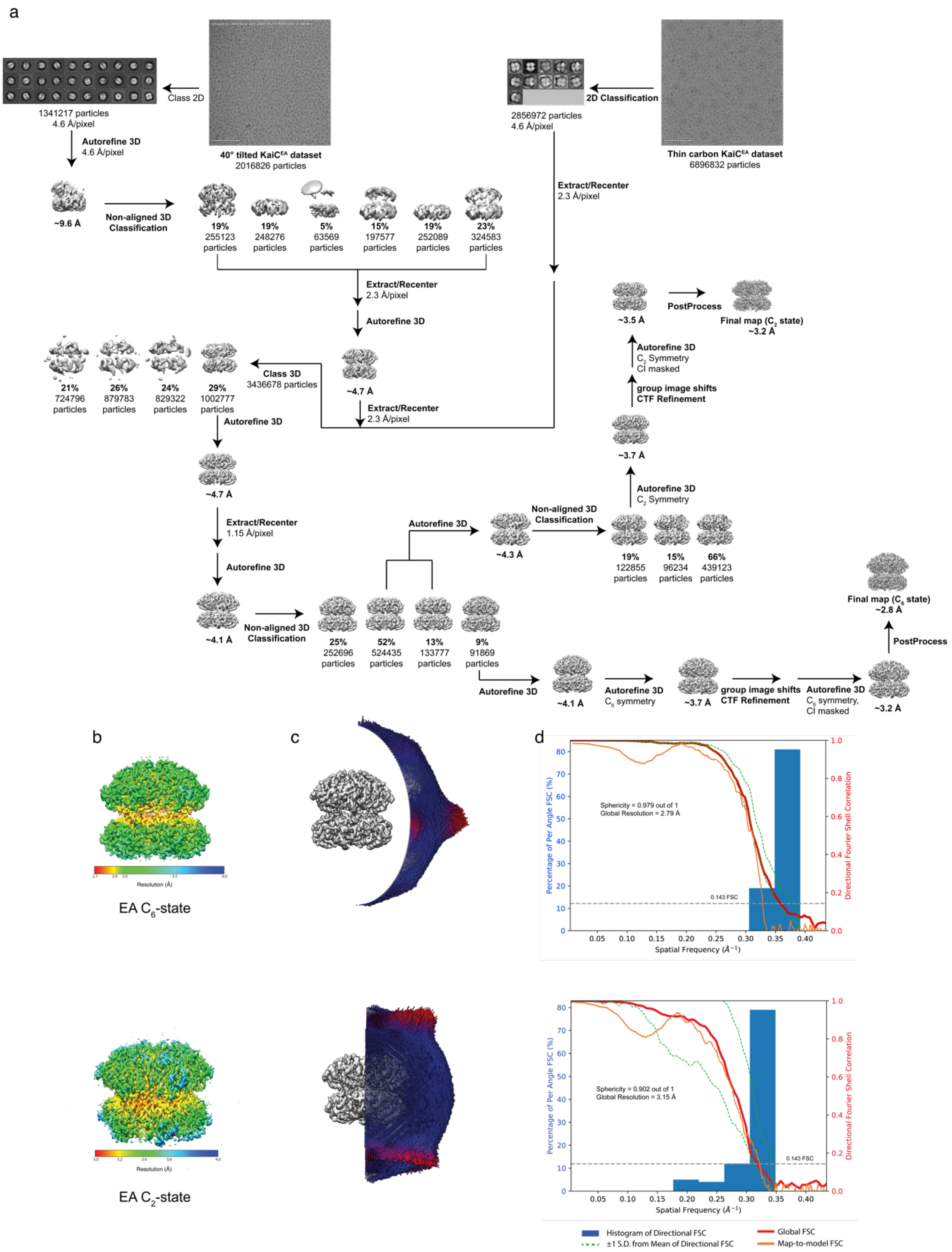


**Fig. S4: Image processing pipeline and validation of KaiC-EA. a**) Image processing pipeline for two combined KaiC-EA datasets: a 40° tilted dataset, and one collected on thin carbon. **b**)

Local resolution estimation of cryo-EM reconstructions calculated by RELION (*33*). **c**) Euler distribution plots depicting particle orientations present in final reconstructions. More populated views are colored in red. **d**) 3D Fourier Shell Correlation (3DFSC) (*34*) of final nighttime state reconstructions, with a global resolution of 2.8 Å for the $C_6$ state, and 3.2 Å for the $C_2$ state at FSC=0.143.

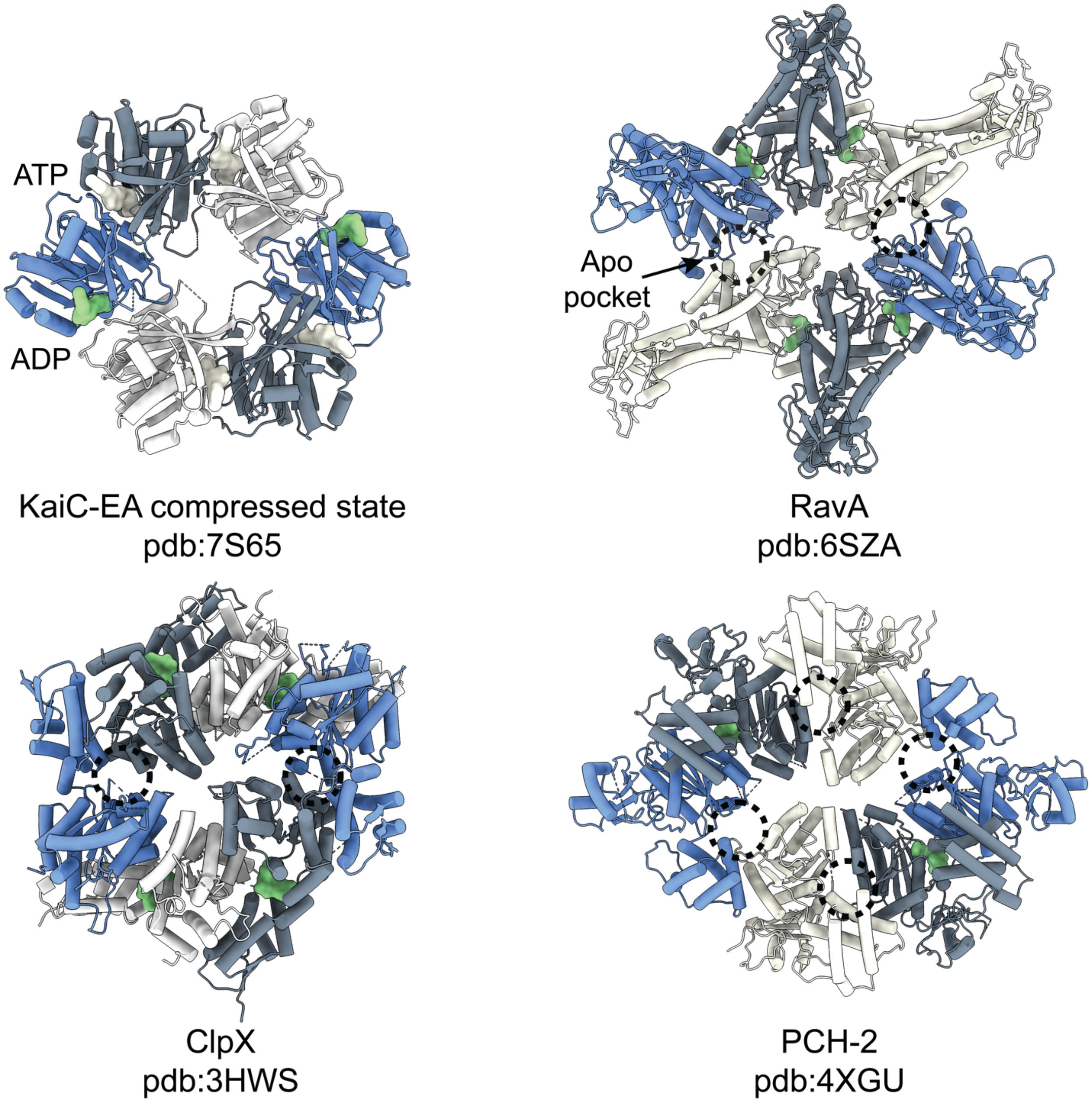


**Figure S5. Structural comparison of $C_2$-symmetric ATPase structures.** Seam protomers are depicted in blue while ATP and ADP nucleotides are shown in white and green, respectively. Apo nucleotide pockets are indicated with dashed circles. While other $C_2$-symmetric ATPases are unliganded at their seam protomers, KaiC is bound to ADP at these subunits.

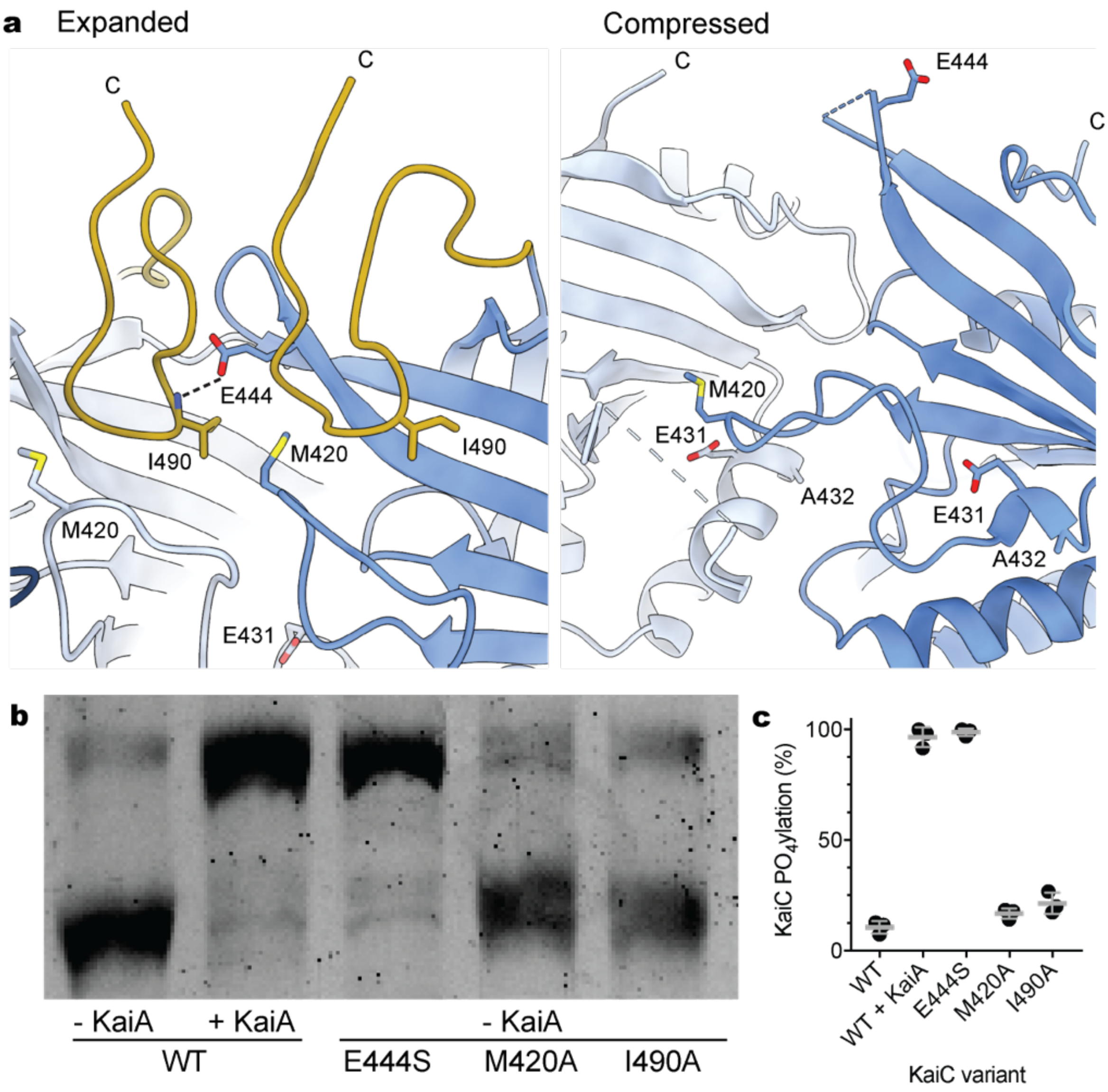


**Figure S6. Allostery about the CII ring regulates KaiC autophosphorylation. a**) In the expanded state, the A-loop of an adjacent protomer (yellow) is positioned near the phosphosite-adjacent 422-loop through a hydrophobic interaction between I490 and M420. The sidechain of E444 forms a hydrogen bond with the mainchain nitrogen of I490 in *trans*. In the compressed state of KaiC-EA, residue E444 is positioned away from the pore in the ADP-bound protomers and no longer stabilizes the A-loops, allowing them to become disordered and causing the 422-loops to collapse towards phosphosite-containing α9. Binding of KaiA to the C-terminus of KaiC may disrupt this tripartite interaction and stimulate hyperphosphorylation. **b**) SYPRO Orange fluorescence of KaiC mutants that were run on a 10% denaturing polyacrylamide gel containing 50μM Phos-tag™ reagent and 100μM $Mn^{2+}$. **c**) Densitometric analysis of bands corresponding to phosphorylated and unphosphorylated KaiC. Gray bars represent mean ± standard deviation for n = 3 repeats.

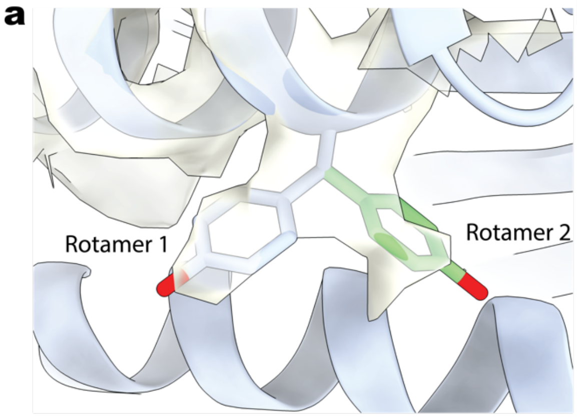


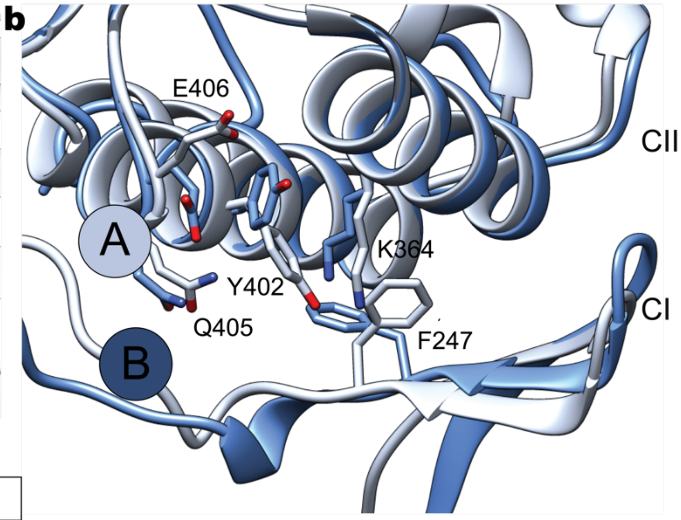


| | Rotamer 1 | Rotamer 2 |
|---|---|---|
| Rotamer probability (Coot) | 43 % | 34 % |
| Residue Model vs. Data CC (Phenix) | 0.74 | 0.65 |

**Figure S7. KaiC residue Y402 exhibits rotameric heterogeneity in the compressed conformation. a**) Close-up view of the two Y402 rotamer conformations from the compressed protomer of the $C_2$-symmetric KaiC-EA structure. Cryo-EM density is depicted in white and semi-transparent for clarity. Analysis from modeling software Coot (*41*) indicates that both rotamers are allowed with similar probabilities. Model vs. Data Cross Correlation in Phenix (*42*) indicates that Rotamer 1 has a higher correlation compared to Rotamer 2, but that both are favorable. **b**) Superposition of compressed protomer B with expanded protomer A, aligned by the CII domain. The rotameric heterogeneity of Y402 in the compressed protomer is associated with rotameric deviations in nearby residues in comparison to an expanded protomer.

**a** — CI

| | | |
|---|---|---|
| S. elongatus | IARYGVEEFVSDNVVILRNVLEGERRRRTLEILKLRGTSHMKGEYPFT | 238 |
| T. elongatus | VARFGVEEFVSDNVVILRNVLEGERRRRTVEILKLRGTTHMKGEYPFT | 239 |
| P. marinus | IARYGVEEFVSDNVVLLRNVLEAEKRRRTLEVLKLRGTVHMKGEFPFT | 234 |
| Nostoc_sp. | VASFGVEEFVSDNVVIARNVLEGERRRRTIEILKLRGTTHMKGEYPFT | 237 |
| A. variabilis | VASFGVEEFVSDNVVIARNVLEGERRRRTIEILKLRGTTHMKGEYPFT | 237 |
| Synechocystis sp. | IARFGVEEFVSDNVVVLRNVLEGERRRRTVEILKLRGTTHMKGEYPFT | 239 |
| M._aeruginosa | VARFGVEEFVSDNVVIMRNVLEGERRRRTAEILKLRGTTHMKGEYPFT | 239 |
| Cyanothece_sp. | IARYGVEEFVSDNVVVLRNVLEGERRRRTAEILKLRGTTHMKGEYPFT | 239 |

**b** — CII

| | | |
|---|---|---|
| S. elongatus | SITDSHISTITDTIILLQYVEIRGEMSRAINVFKMRGSWHDKAIREFM | 471 |
| T. elongatus | SITESHISTITDTILLLQYVEIRGEMSRAINVFKMRGSWHDKGIREYV | 471 |
| P. marinus | SITDSHISTITDTILLLQYVEIKGEMARAINVFKMRGSWHDKRIREYI | 467 |
| Nostoc_sp. | SITDSHISTITDTILMLQYVEIRGEMSRAINVFKMRGSWHDKGIREYN | 470 |
| A. variabilis | SITDSHISTITDTILMLQYVEIRGEMSRAINVFKMRGSWHDKGIREYN | 470 |
| Synechocystis_sp. | SITESHISTITDTILMLQYVEIRGEMSRALNVFKMRGSWHDKGIREYS | 472 |
| M. aeruginosa | SITESHISTITDTILMLQYVEIRGEMSRAINVFKMRGSWHDKGIREYT | 472 |
| Cyanothece_sp. | SITESHISTITDTIIMLQYVEIRGEMSRAINVFKMRGSWHDKGIREYS | 472 |

**Figure S8. Multiple sequence alignment of key KaiC regions from various species of cyanobacteria**. **a**) Protein sequences of the CI and **b**) CII regions of KaiC from eight distinct strains of cyanobacteria obtained from UniProt (*54*) and aligned using Clustal ω (*55*). Key residues are highlighted with colors illustrating their conservation between the CI and CII domains.

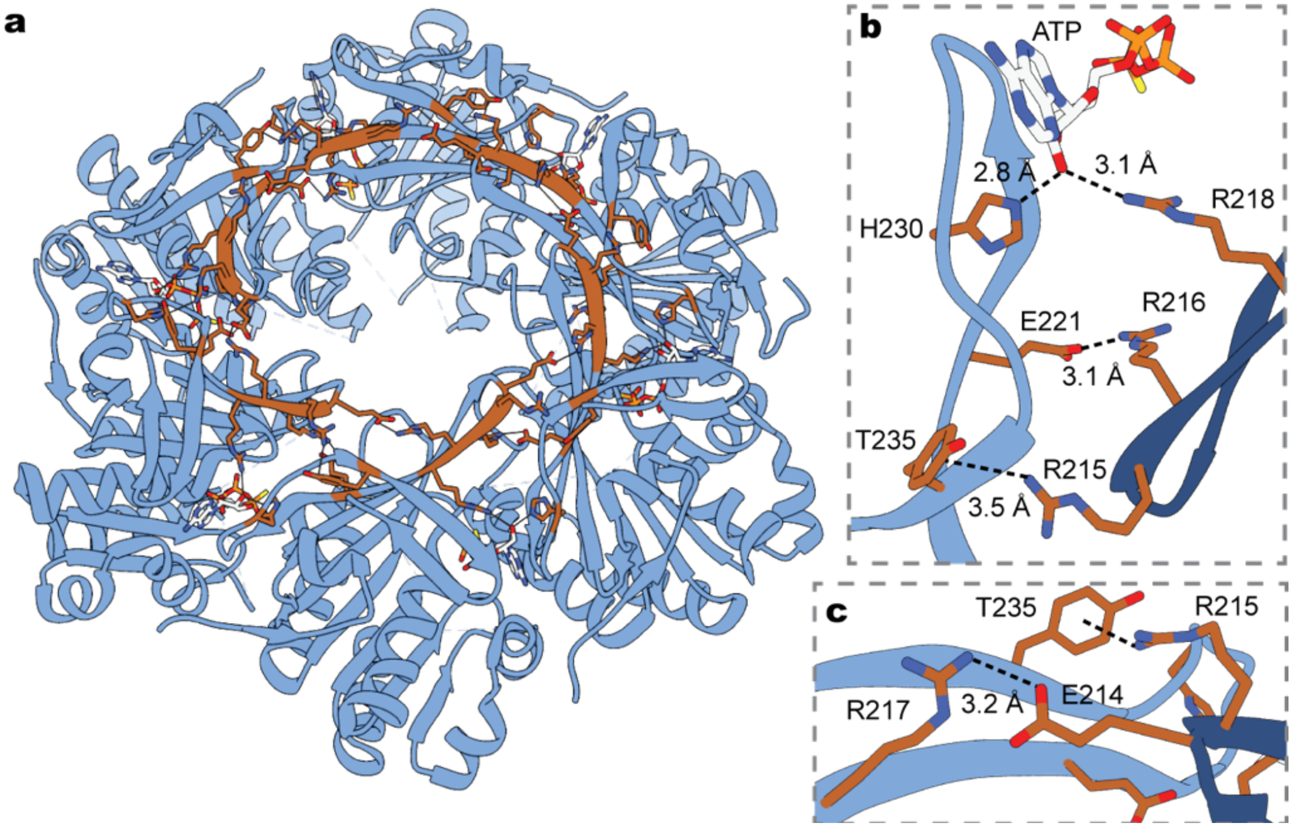


**Figure S9. Interactions of the arginine tetrad create a network of CI-CI *trans* interactions.** **a**) The 4TLA structure of the KaiC CI hexamer is shown as light blue ribbons with the arginine tetrad shown in brown. **b-c**) Close-up views of the arginine tetrad sidechains and their electrostatic interaction partners are shown in brown, with the clockwise protomer in dark blue. The average interatomic distance from the six interfaces are indicated as summarized in Table S1.

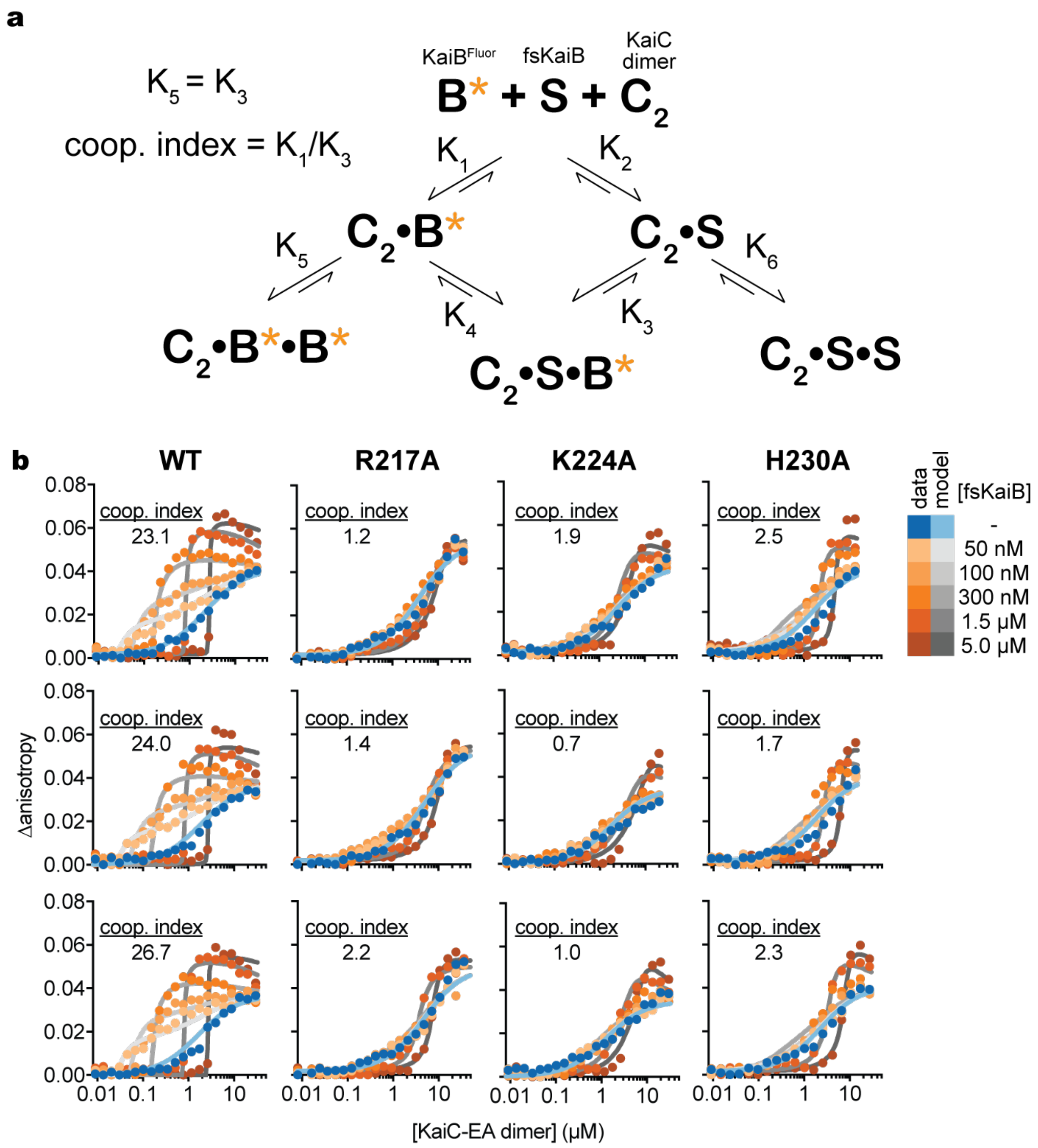


**Figure S10. Cooperativity analysis of KaiC mutants. a**) Thermodynamic model for KaiB homocooperativity, based on a model originally derived for heterotropic cooperativity in (*26*). In this case, the heterotropic cooperativity factor, S, corresponds to unlabeled KaiB-I87A (fsKaiB), which binds much tighter to KaiC than wild-type KaiB (*8*). Binding of wild-type KaiB to KaiC is monitored as indicated by an orange asterisk. **b**) 2-dimensional titrations are overlaid with titration curves derived from least-squares fitting of the data from each individual dataset.

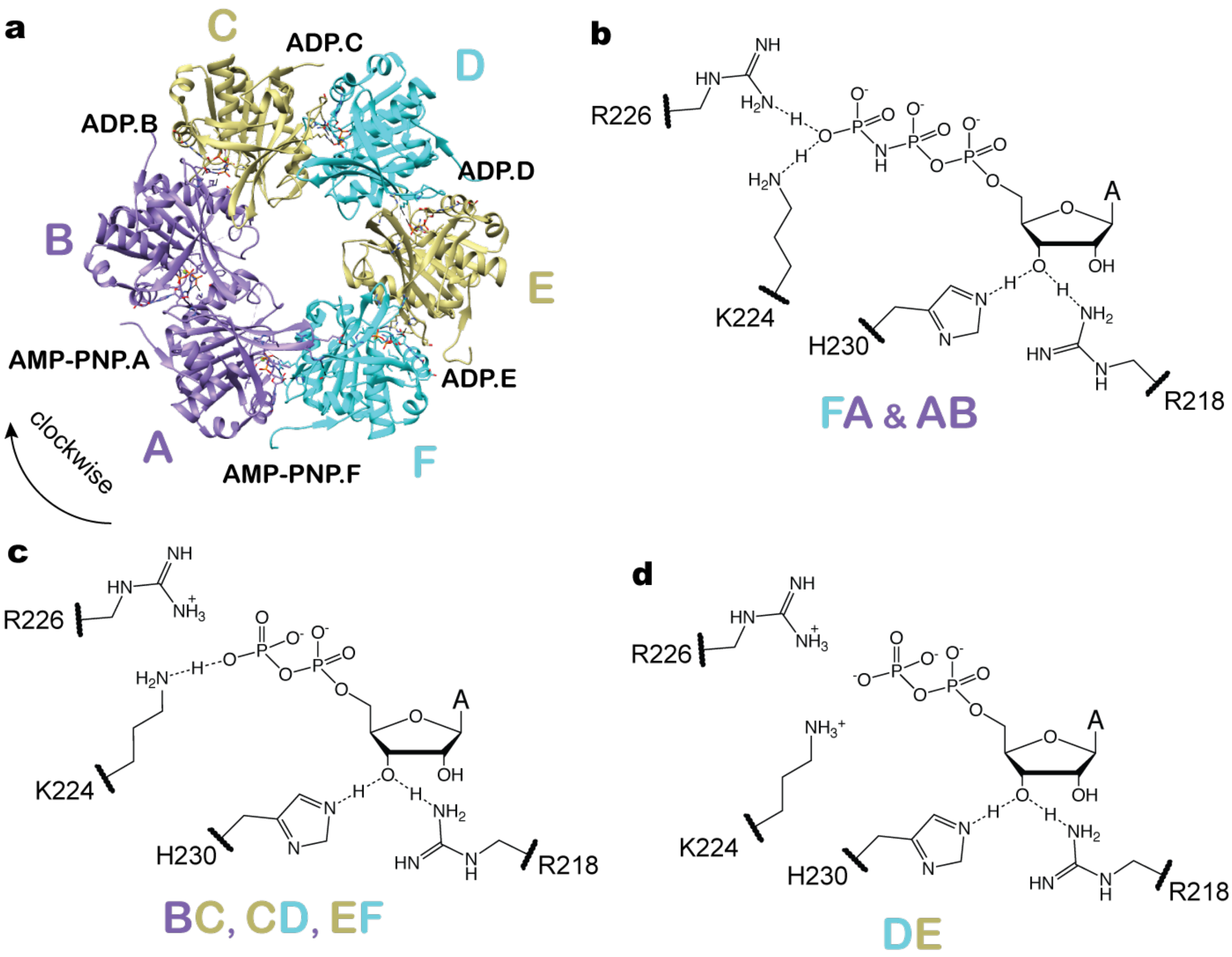


**Figure S11: Analysis of CI-CI nucleotide contacts from PDB 4TLA. a**) Top-down view of the CI domain from the mixed nucleotide state CI structure reported in Abe et al. (*22*) with the nucleotides observed at each interface labeled. Schematic depictions of the specific atomic interactions with the 2’ hydroxyls as well as residues K224 and R226 observed at each interface exhibiting either two (**b**), one (**c**) or no (**d**) electrostatic interactions with the nucleotide phosphates.

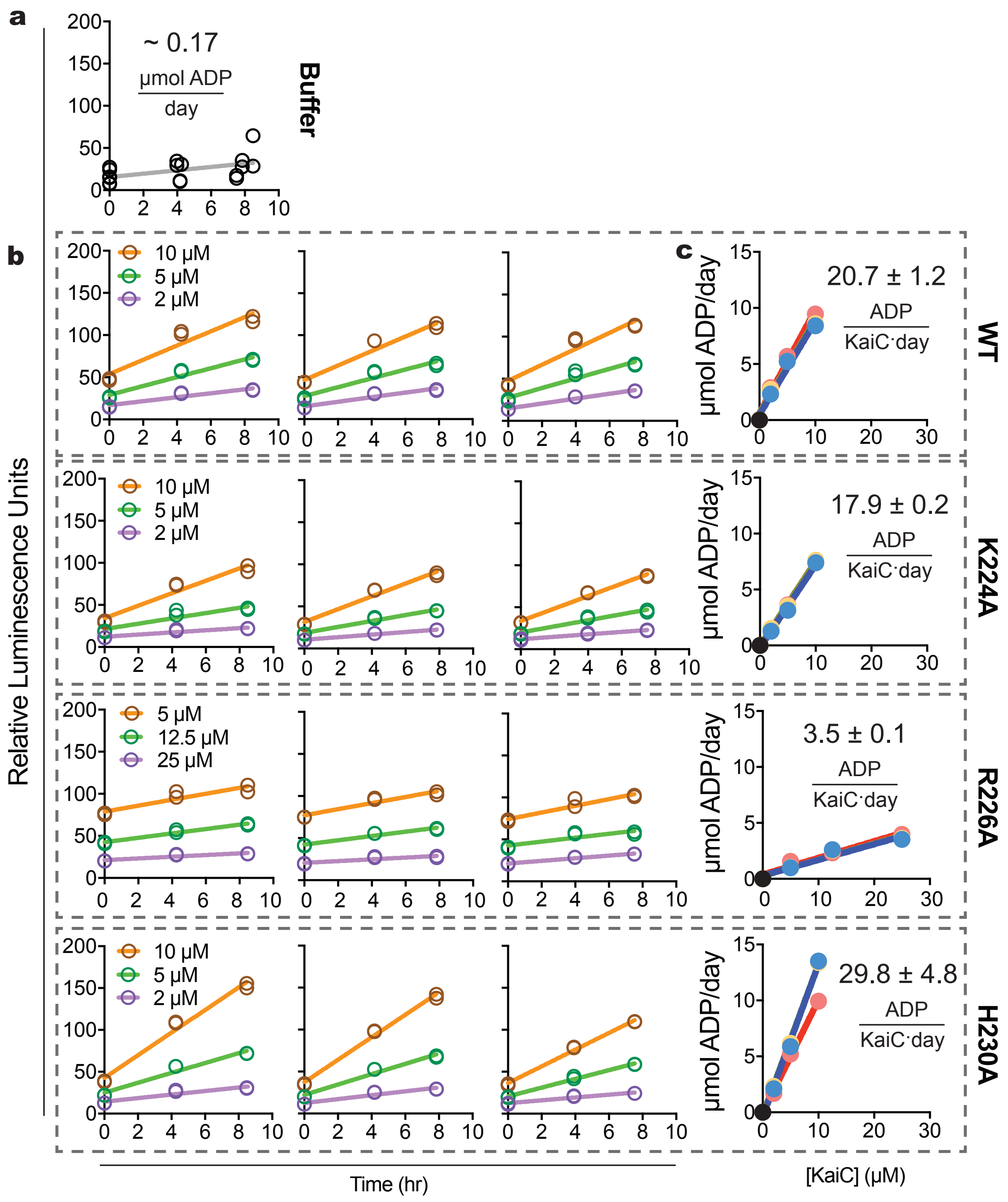


**Figure S12. KaiC ATPase Assays. a**) Triplicate trajectories of luciferase signal in ADP-Glo assays for background ATP hydrolysis from buffer alone, fit to a single regression line giving an overall rate of 0.17 µmol/day. **b**) Luminescence trajectories of KaiC dependent ATP-hydrolysis monitored by ADP-Glo at three concentrations of KaiC-EA. **c**) Turnover rate constants ($k_{cat}$) were subsequently extracted from the slopes of the KaiC trajectories (red, yellow and blue for replicate measurements) plotted against the concentration of the KaiC-EA variant, using the buffer rate as the zero point (black).

**Table S1. Protein constructs and shorthand names used in this study.**

| Shorthand name | Protein construct full name |
|---|---|
| KaiC-AE (daytime KaiC) | FLAG-seKaiC-1-519-S431A-T432E |
| KaiC-EA (nighttime KaiC) | FLAG-seKaiC-1-519-S431E-T432A |
| KaiC-AE-K457A | FLAG-seKaiC-1-519-S431A-T432E-K457A |
| KaiC-EA-K457A | FLAG-seKaiC-1-519-S431E-T432A-K457A |
| KaiC-EA-I430G | FLAG-seKaiC-1-519-I430G-S431E-T432A |
| KaiC-EA-R216A | FLAG-seKaiC-1-519-R216A-S431E-T432A |
| KaiC-EA-R217A | FLAG-seKaiC-1-519-R217A-S431E-T432A |
| KaiC-EA-H230A | FLAG-seKaiC-1-519-H230A-S431E-T432A |
| KaiC-EA-R226A | FLAG-seKaiC-1-519-R226A-S431E-T432A |
| KaiC-EA-K224A | FLAG-seKaiC-1-519-R224A-S431E-T432A |
| KaiC | FLAG-seKaiC-1-519 |
| KaiC-H230A | FLAG-seKaiC-1-519-H230A |
| KaiC-K224A | FLAG-seKaiC-1-519-K224A |
| KaiB-fluorescein | seKaiB-1-102-K25C-FLAG |
| fsKaiB | seKaiB-1-99-Y7A-I87A-Y93A-FLAG |
| KaiA | seKaiA-1-284 |

**Table S2. Cryo-EM data collection, refinement and validation statistics**

| | *KaiC-EA Compressed State* *EMDB: EMD-24850* *PDB: 7S65* | *KaiC-EA Expanded State* *EMDB: EMD-24851* *PDB: 7S66* | *KaiC-AE Expanded State* *EMDB: EMD-24852* *PDB: 7S67* |
|---|---|---|---|
| **Data Collection** | | | |
| Microscope | Talos Arctica | Talos Arctica | Talos Arctica |
| Voltage (KeV) | 200 | 200 | 200 |
| Detector | K2 Summit | K2 Summit | K2 Summit |
| Magnification (nominal/calibrated) | 36,000X/43,478X | 36,000X/43,478X | 36,000X/43,478X |
| Exposure navigation | Image shift to 4 holes (tilted)/ Image shift to 16 holes (thin carbon) | Image shift to 4 holes (tilted)/ Image shift to 16 holes (thin carbon) | Image shift to 16 holes |
| Data acquisition software | Leginon | Leginon | Leginon |
| Total electron exposure ($e^{-}/Å^{2}$) | 48 (tilted)/51 (thin carbon) | 48 (tilted)/51 (thin carbon) | 40.3 |
| Exposure rate ($e^{-}$/pixel/sec) | 7.94 (tilted)/7.5 (thin carbon) | 7.94 (tilted)/7.5 (thin carbon) | 8.6 |
| Frame length (ms) | 100 (tilted)/250 (thin carbon) | 100 (tilted)/250 (thin carbon) | 145 |
| Number of frames per micrograph | 80 (tilted)/ 36 (thin carbon) | 80 (tilted)/ 36 (thin carbon) | 62 |
| Pixel size (Å) | 1.15 | 1.15 | 1.15 |
| Defocus range (μm) | -0.8 to -1.7 (tilted)/ -0.5 to -1.5 (thin carbon) | -0.8 to -1.7 (tilted)/ -0.5 to -1.5 (thin carbon) | -0.5 to -2.0 |
| Micrographs collected (no.) | 1137 (tilted)/8405 (thin carbon) | 1137 (tilted)/8405 (thin carbon) | 1,541 |
| **Reconstruction** | | | |
| Image processing package | Relion | Relion | Relion/CryoSPARC |
| Total extracted particles (no.) | 2016826 (tilted)/ 6896832 (thin carbon) | 2016826 (tilted)/ 6896832 (thin carbon) | 427620 |
| Refined particles (no.) | 1341217 (tilted)/2856972 (thin carbon) | 1341217 (tilted)/2856972 (thin carbon) | 367247 |
| Final particles (no.) | 122855 | 91869 | 89892 |
| Symmetry imposed | C2 | C6 | C6 |
| Global Resolution (Å) | 3.2 | 2.8 | 3.8 |

| | | | |
|---|---|---|---|
| FSC 0.143 (unmasked/masked) | 3.3/3.2 | 2.9/2.9 | 3.9/3.8 |
| FSC 0.5 (unmasked/masked) | 3.5/3.4 | 3.2/3.1 | 4.1/4.0 |
| Resolution range (local) | 3.0 - 4.0 | 2.7 - 4.0 | 3.6 - 4.7 |
| 3DFSC Sphericity | 0.902 out of 1 | 0.979 out of 1 | 0.973 out of 1 |
| Sharpening B-factor (Å$^2$) | -148.4 | -112.9 | -126.7 |
| **Model Composition** | | | |
| Protein residues | 2765 | 2910 | 2892 |
| Ligands | 24 | 24 | 24 |
| **Model Refinement** | | | |
| Refinement package | Phenix | Phenix | Phenix |
| CC (volume/mask) | 0.72/0.74 | 0.81/0.83 | 0.73/0.73 |
| R.m.s deviations | | | |
| Bond lengths | 0.014 | 0.012 | 0.0013 |
| Bond angles | 1.133 | 1.028 | 1.407 |
| **Validation** | | | |
| Map-to-model FSC 0.5 | 3.3 | 3.1 | 3.9 |
| Ramachandran (%) | | | |
| Outliers | 0 | 0 | 0 |
| Allowed | 1.93 | 1.45 | 1.46 |
| Favored | 98.07 | 98.55 | 98.54 |
| MolProbity score | 1.4 | 0.99 | 1.28 |
| Poor rotamers (%) | 0.17 | 0 | 0 |
| Clashscore (all atoms) | 8.20 | 2.16 | 5.19 |
| C-beta deviations (%) | 0 | 0 | 0.23 |
| CaBLAM Outliers (%) | 0.8 | 1 | 0.8 |
| EMRinger Score | 3.52 | 3.8 | 2.1 |

**Table S3. Interatomic distances for *trans* interactions of the arginine tetrad.** Distances measured between the specified atoms at each of the 6 interfaces from PDB 4TL8.

| **Interface** | **Interatomic Distances (Å) from 4TL8** | | | | |
|---|---|---|---|---|---|
| **AB** | R217.B NH - E214.A<br>3.1 | R216.A NH - E221.B Oε<br>2.9 | R215.A NH - Y235.B Cζ<br>3.8 | R218.A NH - AGS.A O2'<br>3.0 | H230.B Nε - AGS.A O2'<br>2.8 |
| **BC** | R217.C NH - E214.B<br>3.0 | R216.B NH - E221.C Oε<br>2.9 | R215.B NH - Y235.C Cζ<br>3.6 | R218.B NH - AGS.B O2'<br>3.1 | H230.C Nε - AGS.B O2'<br>2.8 |
| **CD** | R217.D NH - E214.C<br>4.2 | R216.C NH - E221.D Oε<br>3.3 | R215.C NH - Y235.D Cζ<br>3.3 | R218.C NH - AGS.C O2'<br>3.0 | H230.D Nε. AGS.C O2'<br>2.8 |
| **DE** | R217.E NH - E214.D<br>3.1 | R216.D NH - E221.E Oε<br>3.4 | R215.D NH - Y235.E Cζ<br>3.3 | R218.D NH - AGS.D O2'<br>3.0 | H230.E Nε AGS.D O2'<br>2.8 |
| **EF** | R217.F NH - E214.E<br>2.9 | R216.E NH - E221.F Oε<br>3.2 | R215.E NH - Y235.F Cζ<br>3.5 | R218.E NH AGS.E O2'<br>3.2 | H230.F Nε AGS.E O2'<br>2.8 |
| **FA** | R217.A NH - E214.F<br>3.0 | R216.F NH - E221.A Oε<br>3.0 | R215.F NH - Y235.A Cζ<br>3.6 | R218.F NH - AGS.F O2'<br>3.1 | H230.A Nε AGS.F O2'<br>2.8 |

**Table S4. Pre and post-hydrolysis distances in active site interactions of the CI domain.** Distances measured between the specified atoms in each of the 6 interfaces from PDB 4TLA (*22*).

| **Interface** | **Interatomic Distances (Å) from 4TLA** | | | |
|---|---|---|---|---|
| **AB** | K224.B Nζ - ANP.A Oγ<br>2.6 | R226.B NH - ANP.A Oγ<br>2.6 | H230.B Nε - ANP.A O2'<br>2.8 | R218.A NH - ANP.A O2'<br>3.0 |
| **BC** | K224.C Nζ - ADP.B Oβ<br>2.5 | R226.C NH - ADP.B Oβ<br>5.0 | H230.C Nε - ADP.B O2'<br>2.8 | R218.B NH - ADP.B O2'<br>3.0 |
| **CD** | K224.D Nζ - ADP.C Oβ<br>2.7 | R226.D NH - ADP.C Oβ<br>5.0 | H230.D Nε. ADP.C O2'<br>2.9 | R218.C NH - ADP.C O2'<br>3.0 |
| **DE** | K224.E Nζ - ADP.D Oβ<br>4.4 | R226.E NH ADP.D Oβ<br>4.9 | H230.E Nε ADP.D O2'<br>2.8 | R218.D NH - ADP.D O2'<br>3.1 |
| **EF** | K224.F Nζ - ADP.E Oβ<br>2.6 | R226.F NH - ADP.E Oβ<br>5.6 | H230.F Nε ADP.E O2'<br>2.7 | R218.E NH ADP.E O2'<br>3.2 |
| **FA** | K224.A Nζ - ANP.F Oγ<br>2.8 | R226.A NH - ANP.F Oγ<br>2.9 | H230.A Nε ANP.F O2'<br>2.8 | R218.F NH - ANP.F O2'<br>3.1 |

**Table S5. Plasmids and primers used in generating cyanobacterial strains**

| Plasmids | Description | Source |
|---|---|---|
| pSL2680 | CRISPR/Cas12a plasmid; Km resistance | Addgene (#85581) |
| pSL2680-K224A | pSL2680 + KaiC K224A substitution | This study |
| pSL2680 | RSF1010-backbone, Cas12a and CRISPR assay from *Francisella novicida*, $Km^R$ | Ungerer and Pakrasi, 2016 (*9*) |
| **Primers** | **Sequence (5'-3')** | |
| K224A gRNA F | AGATgaggatttcgagggtgcggc | |
| K224A gRNA R | AGACgccgcaccctcgaaatcctc | |
| K224A homology arm upstream F | ggtgccacgtagcGCgaggatttcgagggtg | |
| K224A homology arm upstream R | gcccggattacagatcctctagagtcgacGGTACCCCTCGCCAAGGTTCTACCAC | |
| K224 homology arm downstream R | catttttttgtctagctttaatgcggtagttGGTACCTCGACGGTAATCCGTGTTGG | |
| K224A homology arm downstream R | caccctcgaaatcctcGCgctacgtggcacc | |

**Table S6. Cyanobacterial strains used in this study**

| Strain | Genotype (NS denotes neutral site) | Antibiotic resistance | Source |
|---|---|---|---|
| WT (AMC541) | NSII-P$_{kaiBC}$::*luc* | Cm | Lab collection |
| Δ*kaiC* (AMC AMC704) | NSII-P$_{kaiBC}$::*luc* Δ*kaiC* | Cm | Lab collection |
| *KaiC*-K224A | NSII-P$_{kaiBC}$::*luc*. *kaiC*-K224A | Cm | This study |

**Data S1. Raw and background subtract4ed data from KaiC mutant titrations (separate file).** Summary of data from Fig. S1 organized by KaiC mutant

**Data S2. DynaFit script for modeling of 2D titration assays (separate file).** Text file containing annotated DynaFit script for two-site binding model used to for cooperativity modeling.

**Data S3. Raw and background subtracted data from 2D titrations and cooperativity modeling of KaiC mutants (separate file).** Summary of raw data, thermodynamic models and curve fitting parameters from Fig. S9 organized by KaiC mutant.
